# Evaluating models of the ageing BOLD response

**DOI:** 10.1101/2023.08.24.554634

**Authors:** R.N. Henson, W. Olszowy, K.A. Tsvetanov, P.S. Yadav, Cam-CAN, P. Zeidman

## Abstract

Neural activity cannot be directly observed using fMRI; rather it must be inferred from the hemodynamic responses that neural activity causes. Solving this inverse problem is made possible through the use of forward models, which generate predicted hemodynamic responses given hypothesised underlying neural activity. Commonly-used hemodynamic models were developed to explain data from healthy young participants; however studies of ageing and dementia are increasingly shifting the focus towards elderly populations. We evaluated the validity of a range of hemodynamic models across the healthy adult lifespan: from basis sets for the linear convolution models commonly used to analyse fMRI studies, to more advanced models including nonlinear fitting of a parameterised hemodynamic response function (HRF) and nonlinear fitting of a biophysical generative model (hemodynamic modelling, HDM). Using an exceptionally large sample of participants, and a sensorimotor task optimized for detecting the shape of the BOLD response to brief stimulation, we first characterised the effects of age on descriptive features of the response (e.g., peak amplitude and latency). We then compared these to features from more complex nonlinear models, fit to four regions of interest engaged by the task, namely left auditory cortex, bilateral visual cortex, left (contralateral) motor cortex and right (ipsilateral) motor cortex. Finally, we validated the extent to which parameter estimates from these models have predictive validity, in terms of how well they predict age in cross-validated multiple regression. We conclude that age-related differences in the BOLD response can be captured effectively by models with three free parameters. Furthermore, we show that biophysical models like the HDM have predictive validity comparable to more common models, while additionally providing insights into underlying mechanisms, which go beyond descriptive features like peak amplitude or latency, and include estimation of nonlinear effects. Here, the HDM revealed that most of the effects of age on the BOLD response could be explained by an increased rate of vasoactive signal decay and decreased transit rate of blood, rather than changes in neural activity per se. However, in the absence of other types of neural/hemodynamic data, unique interpretation of HDM parameters is difficult from fMRI data alone, and some brain regions in some tasks (e.g, ipsilateral motor cortex) can show responses that are more difficult to capture using current models.

## Introduction

Functional magnetic resonance imaging (fMRI) is often used to infer changes in neural activity across the brain as a function of a task or the individual being scanned. However, the Blood Oxygenation Level Dependent (BOLD) signal measured in most fMRI experiments is a complex function of neural activity, neurovascular coupling, blood flow, blood oxygenation and blood volume (Buxton et al., 1998). Therefore, differences in the BOLD response across individuals can be caused by differences in hemodynamics, rather than in neural activity per se. Here we focus on the effects of age, since it is likely that age affects vascular factors as well as neural activity (Handwerker et al., 2012; Tsvetanov et al., 2021; Wright & Wise, 2018). Indeed, neurovascular ageing is not only a confound in fMRI, but an important process to assess alongside neural ageing (Abdelkarim et al., 2019; Tsvetanov et al., 2021).

The most common way to model the BOLD impulse response to an experimental input (e.g., stimulus or trial, assumed to cause a short burst of neural activity) is with a linear convolution model, in which a delta function is convolved with a basis set of temporal functions, typically lasting 0-30s post-stimulus. After convolution with multiple, sequential inputs in a typical fMRI experiment, each convolved basis function forms a separate regressor within a general linear model (GLM). When fitting the GLM to the fMRI timeseries from a voxel or region of interest (ROI), the weights (parameter estimates) for each such function can be estimated using ordinary least squares minimisation. There are multiple options for such basis sets (Henson, 2004), but a common one is the Finite Impulse Response (FIR) basis set, which consists of multiple “top-hat” functions, e.g., 32 contiguous time bins of 1 s duration, spanning from 0 to 32 s post-stimulus (Figure 1).

**Figure 1.**
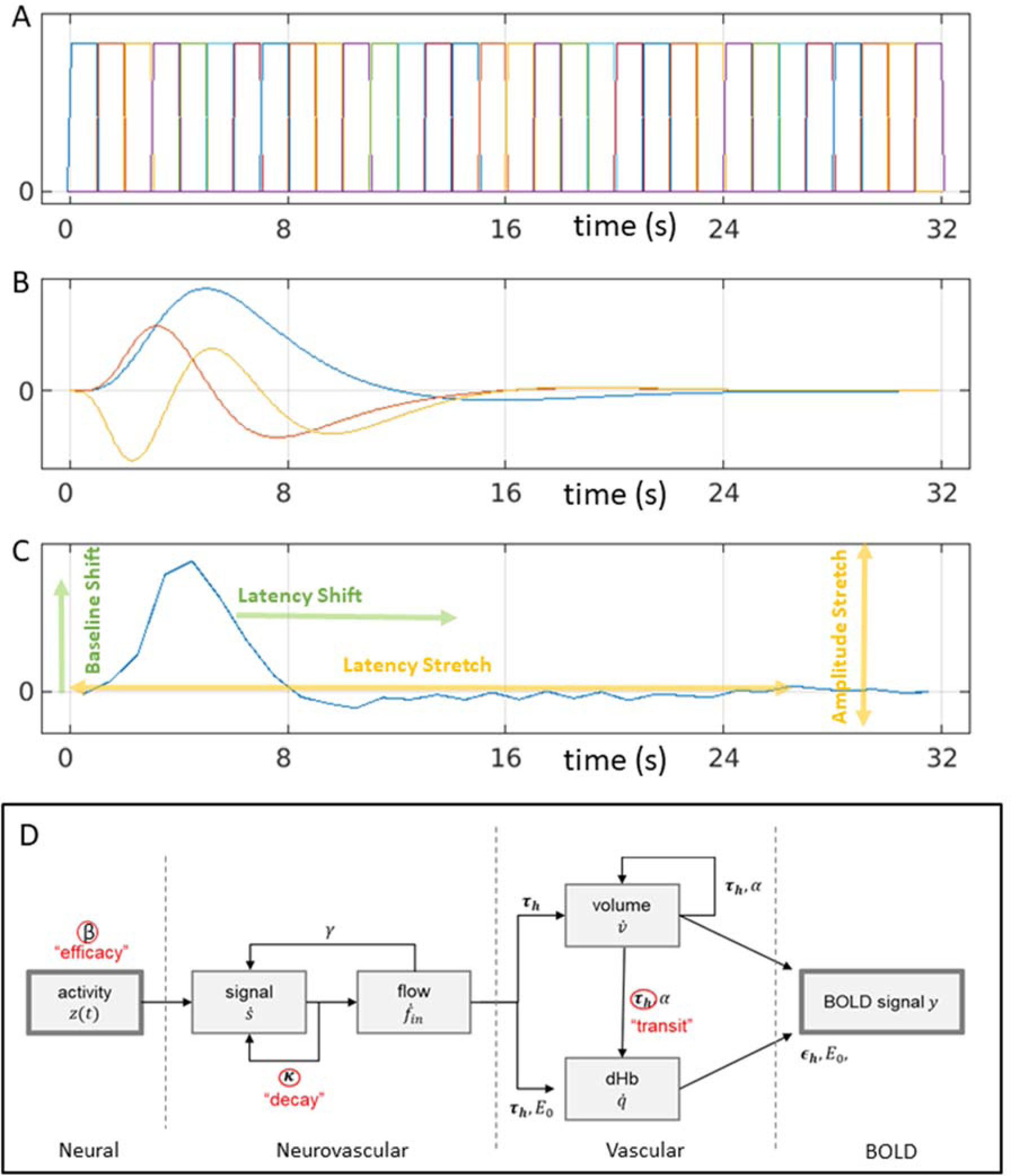
HRF models: (A) linear Finite Impulse Response basis set with 32 x 1s bins (FIR32); (B) an “informed” linear basis set with a canonical HRF (blue), its temporal derivative (red) and dispersion derivative (yellow) (Can3); (C) nonlinear fitting of average empirical HRF for a given ROI (here lAC) using amplitude and latency offsets and scalings (NLF4); (D) nonlinear hemodynamic modelling using differential equations with three free parameters (in red) (HDM3).

Using such a flexible basis set (whether the FIR set in the time domain, or sinusoidal functions of periodic frequencies in the Fourier domain), the first study to address age effects failed to find differences in BOLD response shape between young versus older participants (D’Esposito et al., 1999). However, like many studies, this study used relatively small groups (n=32 young and n=20 older), and the older volunteers may have been healthier than average for their age. Furthermore, tests using such flexible basis sets may be less sensitive to subtle statistical differences in HRF shape.

Even though the BOLD response can vary across individuals and brain regions, it has a typical shape: peaking around 5s and with an undershoot lasting 10-30s: often called the canonical hemodynamic response function (HRF; see Figure 1 for an example). Thus a common approach is to assume this canonical form, and allow for some variability around this by adding, e.g, its partial derivatives in SPM’s “informed basis set”, or the first few singular vectors in FSL’s data-driven FLOBS approach (Woolrich et al., 2004; see Lindquist et al., 2009, for yet further models of HRF shape). Greater sensitivity to ageing may be achieved by focusing on specific features of the HRF shape, such as the latency of its peak. For example, Taoka et al. (1998) and Handwerker et al. (2007) found an age-related delay in the time to peak, while Richter & Richter (2003) and Aizenstein et al. (2004) found age-related reductions in the magnitude of the undershoot. By contrast, other studies have reported no effect of age. For example, Ward et al. (2008) found no age effects on the HRF in contralateral motor cortex during hand grips, while Grinband et al. (2017) found few effects of age on various HRF features, and argued that those found previously could reflect age-related differences in neural activity. Moreover, even if differences are found in the shape of the HRF (e.g, peak latency), relating these to differences in underlying neural activity is far from simple, given that the mapping from neural activity to BOLD response is nonlinear (see Discussion).

Again, the above studies involved relatively small samples. In the largest study of HRF shape of which we are aware, West et al. (2019) compared a younger (n=74) and an older (n=173) group. They found that HRFs differed between groups in auditory, visual and motor cortices, with increased time-to-peak and decreased peak amplitude in older compared to younger adults. These groups are actually a sub-set of the full set of n=645 participants used here. These participants come from the CamCAN cohort (www.cam-can.org), approximately uniformly distributed across 18-88 years of age. These participants were recruited using an opt-out procedure after contact from local surgeries, making them more representative of healthy adults than most typical studies that use opt-in recruiting via adverts (though the present participants are still biased towards people able to lie in an MRI scanner for an hour, and towards those without physical or mental health problems). More specifically, these data come from the CamCAN “sensorimotor” task, in which participants saw brief trials consisting of bilateral visual checkerboards and/or auditory tones (Shafto et al., 2014). Importantly, the time between trials (in terms of the stimulus onset asynchrony, SOA) was deliberately optimised for detecting the shape of the HRF (assuming linear superposition of overlapping responses), by using a 255-length “m-sequence” (Buračas & Boynton, 2002). Here we extend the work of West et al. (2019) by examining the parametric effect of age across the whole sample, and more importantly, by directly comparing different linear and nonlinear methods to characterise age-related changes in the BOLD response.

This study addresses four objectives:

1. To characterise the effects of age on the BOLD response, using a model that makes minimal assumptions about the form of that response, at the cost of a large number of unconstrained parameters (a 32-parameter Finite Impulse Response model, FIR32) used as a basis set within the GLM.
2. To compare these effects of age against those inferred using a more parsimonious basis set with fewer parameters, which is statistically more efficient, but makes stronger assumptions: specifically, the 3-parameter, canonical or “informed” basis set used in the SPM software package (Can3).
3. To compare these linear fits to a potentially more flexible, 4-parameter model, in which the parameters refer to specific features of the BOLD response shape, such as latency and amplitude, and which are non-linearly fit to the trial-averaged FIR estimates (a Non-Linear Fitting approach, NLF4).
4. To go beyond the descriptive statistics provided by the models above, and fit a mechanistic, biophysical Hemodynamic Model (HDM) to the same data. This model explains the effects of ageing in terms of 3 biologically-meaningful parameters: neural efficacy, neurovascular signal decay rate and hemodynamic transit rate. A version of this HDM is widely used as part of the Dynamic Causal Modelling (DCM) framework used for connectivity analysis (Friston et al., 2003).
5. To test whether the models’ parameters have real-world (predictive) validity. We did this by evaluating how well all the above models can predict the age of previously unseen participants, using cross-validation, and whether they relate to independent measures of vascular health and neural activity.

## Methods

### Participants

The N=645 participants were aged 18-88, fluent English speakers and in good physical and mental health based on the CamCAN cohort exclusion criteria, which excluded volunteers with a low Mini Mental State Examination score (<24), serious current medical or psychiatric problems or poor hearing or vision, and standard MRI contraindications (for more details, see Shafto et al., 2014). None showed evidence of neurodegeneration in their structural MRIs (though we cannot rule out sub-clinical effects of neural and/or vascular disease that increase with age). The study was conducted in compliance with the Helsinki Declaration. All participants gave written informed consent for the study and record linkage, as approved by the local ethics committee, Cambridgeshire 2 Research Ethics Committee (reference: 10/H0308/50). Nine participants were removed because they did not show a significant evoked response in at least one of the four ROIs (see ROI section below), leaving 636 participants, exactly half (318) of each sex.

### Paradigm

Each trial consisted of two oval checkerboards containing a rectangular pattern presented either side of a central fixation cross (34 ms duration) and/or a simultaneously-onsetting binaural auditory tone (300 ms duration). The inter-stimulus interval was a blank screen containing only the fixation cross (to which participants were told to fixate throughout). The auditory tones were one of three equiprobable frequencies (300 Hz, 600 Hz, or 1200 Hz). Frequency was not relevant to the task or current hypotheses, so is collapsed here. The majority of the trials (120 of the 128) involved both visual and auditory stimuli, which required a keypress with the right index finger; the remaining 8 randomly-intermixed “catch” trials had only visual (4 trials) or only auditory (4 trials) stimuli, and participants were told to withhold their responses to such “unimodal” trials. The latter ensured that participants needed to pay attention to both modalities in all trials. Visual acuity was corrected with MRI-compatible glasses, while auditory presentation intensity was adjusted so that the tones were audible for all participants.

The stimulus onset asynchrony (SOA) between trials was determined by a second-order m-sequence (m = 2; Buračas & Boynton, 2002), in which half of the sequence elements were null events. The minimal SOA was 2 s, with the null events serving only to produce SOAs that ranged from 2 to 26 s (null events do not produce any change in stimulation when they occur). There was an additional random jitter drawn from a uniform distribution of 0.1-0.3s before stimulus onset, which meant that the effective sampling interval of the HRF across trials was 0.12s (Josephs & Henson, 1999). The total duration of the task was 8mins, 34secs (n=261 volumes).

### MRI data acquisition

The MRI data were collected using a Siemens Trio 3T MRI Scanner system with a 32-channel head coil. A T2*-weighted echo planar imaging (EPI) sequence was used to collect 261 volumes, each containing 32 axial slices (acquired in descending order) with slice thickness of 3.7 mm and an interslice gap of 20% (for whole-brain coverage including cerebellum; repetition time = 1970 ms; echo time = 30 ms; flip angle = 78°; field of view = 192 mm x 192 mm; voxel size 3 x 3 x 4.44 mm). Higher resolution (1 x 1 x 1 mm) T1- and T2-weighted structural images were also acquired to aid registration across participants. For more details, see https://camcan-archive.mrc-cbu.cam.ac.uk/dataaccess/pdfs/CAMCAN700_MR_params.pdf.

### MRI data preprocessing

MRI preprocessing used the SPM12 software (Wellcome Centre for Human Neuroimaging; https://www.fil.ion.ucl.ac.uk/spm), release 4537, implemented in the Automatic Analysis pipeline, release 4.2 (Cusack et al., 2015). Preprocessing is described in (Taylor et al., 2017), but in brief, structural images were rigid-body registered to a Montreal Neurological Institute (MNI) template brain, bias corrected, segmented, and warped to match a gray matter template created from the whole CamCAN Stage 2 sample using SPM’s DARTEL toolbox. This template was subsequently affine transformed to standard MNI space. The functional images were spatially realigned, interpolated in time to the middle slice to correct for the different slice acquisition times, rigid-body coregistered to the structural image, transformed to MNI space using the warps and affine transforms from the structural image, and resliced to 3 mm x 3 mm x 3 mm voxels. An XML summary of preprocessing can be found here: https://camcan-archive.mrc-cbu.cam.ac.uk/dataaccess/ImagingScripts/mri_aa_release004_roistreams_v1_tasklist.xml.

### Single-participant (1^st^-level) models

Two models were run, where neural activity was assumed to be: 1) locked to the onset of each audiovisual stimulus or 2) locked to the subsequent key-press. We included unimodal trials for the stimulus-locked model (since they still involved stimulation), but only included them for the response-locked model if the participant incorrectly pressed the key.

GLMs for each participant were created and estimated in SPM release 7771^1^ in Matlab R2020b (https://www.mathworks.com). A single vector of delta functions at the onset of each trial was convolved with each of the temporal basis functions (FIR or SPM’s informed basis set; see below) in a high-resolution space with 32 time-points every TR (resulting in a microtime resolution of 0.062s), and subsequently down-sampled at the middle point (to match the reference slice for the slice-timing correction). Six additional regressors, representing the three rigid body translations and the three rotations estimated in the realignment stage, were included to capture residual movement-related artifacts, plus the timeseries extracted for WM and CSF compartments, to further remove non-neural noise. Finally, the data were scaled to a grand mean of 100 over all voxels and scans, and the model fit to the data in each voxel. The autocorrelation of the error was modelled using a “first-order autoregressive plus white-noise” model, together with a set of cosines that were used to high-pass filter both the model and the data at a frequency cutoff of 1/128 Hz. The GLM and noise model were estimated using restricted maximum likelihood (ReML), and the resulting estimate of error autocorrelation used to prewhiten both the model and the data. Finally, OLS was used to estimate the model parameters for the whitened data.

### Group model for FIR basis set

For the FIR basis set, the parameter estimate for each of the 32 x 1s time bins for each participant was entered into a second-level GLM (equivalent to a repeated-measures ANOVA), as well as a linear and a quadratic modulation by each FIR parameter by age (see Figure 2). Analyses were restricted to a gray-matter mask, determined by thresholding the DARTEL template for voxels with >50% proportion of gray-matter.

**Figure 2.**
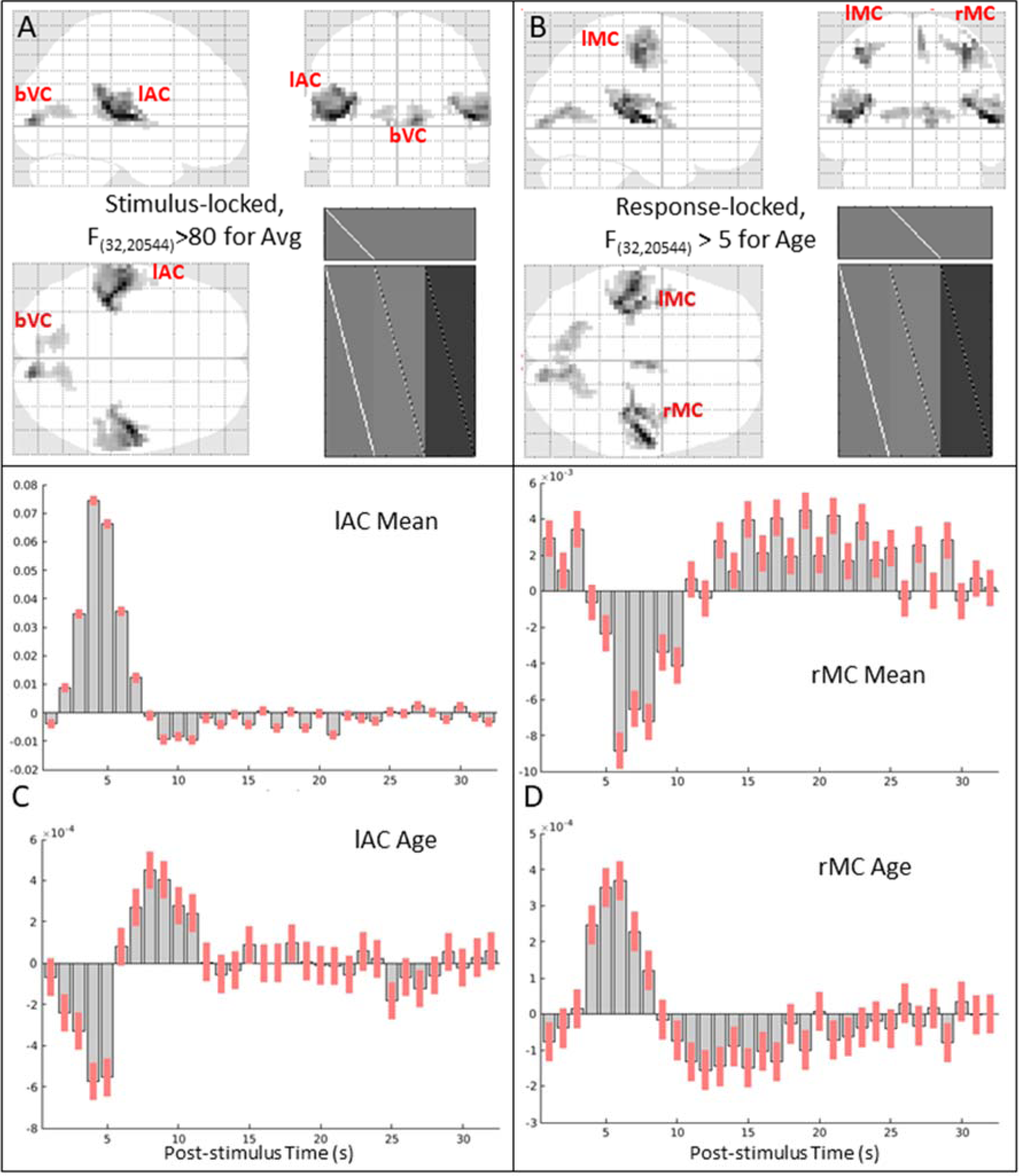
Panels A and B show maximal intensity projections (MIPs) of statistical parametric maps (SPMs) of F-contrast for (A) the mean effect across participants, locked to audio-visual stimulation (thresholded for 50 contiguous voxels with F>80), and (B) the (linear) effect of age, locked to the right finger press (thresholded for 50 contiguous voxels with F>5). The clusters corresponding to the four functional ROIs analysed below (lAC, bVC, lMC and rMC) are labelled on the sagittal, coronal and transverse sections, along with the F-contrast and the design matrix (bottom right). Panels C and D show the mean of each FIR parameter (upper plot) and effect of age on each parameter (lower plot) for the peak voxel from the (C) lAC ROI [-39 -33 +12] and (D) rMC ROI [+36 -21 +51]. Note that the scale of the y-axis is in arbitrary units and differs across plots; for % signal change, see Figure 4. Red bars show 90% confidence interval.

### ROI definition

To maximise sensitivity to age, while not making assumptions about the shape of the BOLD response, masks for each ROI were defined from an F-contrast in the second-level group FIR analysis that spanned the (linear) effects of age.^2^ For the left auditory cortex (lAC) and bilateral visual cortex (bVC), these came from the stimulus-locked model, while for the left and right motor cortex (lMC and rMC), these came from the response-locked model. The F-values were thresholded at F>5 to define clusters of contiguous voxels^3^. This resulted in a mask of 280 and 182 voxels for lAC and bVC, respectively, and 54 and 88 voxels for lMC and rMC, respectively.

To allow for variability across participants in their most responsive voxels, we sub-selected those voxels within the above masks that showed a significant F-contrast (at p<.05 uncorrected) across time bins within each participant’s first-level FIR models. Note that this entailed removing 9 participants with no voxels that survived this threshold. These 9 were roughly equally distributed across age (22, 37, 41, 51, 52, 57, 64, 71, 80 years) and showed excessive motion artifacts in their data (that rigid-body realignment could not correct). The median number of voxels across the remaining participants that passed this threshold was 134, 77, 34 and 21 for lAC, bVC, lMC and rMC respectively. Spearman correlations showed that this number did decrease significantly with age for all ROIs, but we believe that matching the minimal signal-to-noise ratio across age is more important than matching the number of voxels.

The fMRI timeseries for each ROI were then summarised in terms of the first temporal component from a singular-value decomposition (SVD) across the voxels remaining from the above F-contrast, using standard tools for ROI extraction in SPM. This entailed the high-pass filtering, pre-whitening and correction for confounds described above during GLM estimation.

### HRF models

For each ROI, we fit 4 types of model: two linear models (estimated using OLS) and two nonlinear models (fit by gradient descent). The two linear models corresponded to 1) the 32 x 1s bin FIR basis set described above (and used to define the ROIs) – henceforth the “FIR32” model – and 2) SPM’s informed basis set, consisting of a canonical HRF and two partial derivatives – henceforth the “Can3” model. The two nonlinear models were based on shape-matching of the HRF using 4 parameters – the “NLF4” model – and a biophysical generative model with 3 parameters – the “HDM3” model. These are detailed below.

#### FIR32 model

The “FIR32” model consisted of 32 top-hat temporal basis functions of 1s duration that captured the first 32 seconds of post-stimulus time (Figure 1A). This FIR basis set is flexible, but can also capture non-hemodynamic trial-locked effects (e.g, motion of the eyeballs during first time bin, in response to a visual stimulus).^4^ This approach is also called “selective averaging” (Dale & Buckner, 1997).

#### Can3 model

The canonical HRF used in SPM is a mixture of two gamma functions, one capturing the positive peak around 5s and one capturing the negative undershoot peaking around 15s. The parameters of these gamma functions were estimated from approximating the first singular vector of stimulus-locked responses in a previous fMRI experiment (Friston et al., 1998). The second basis function in this “informed” basis set is the temporal derivative of the canonical HRF, obtained by the finite difference after shifting the onset of the canonical HRF by 1 second. The third basis function is the dispersion derivative, obtained by the finite difference after increasing the dispersion of the gamma function for the peak response by 1% (see Figure 1B).

#### NLF4 model

The NLF4 model refers to nonlinear fitting of a template HRF to each individual’s FIR fit, using a first order expansion with respect to amplitude and with respect to time (i.e., four parameters in total). More precisely, the following model is fit:

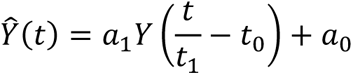

where vector *Y* is the template HRF, derived from the first singular vector from an SVD of the FIR values across all participants, *t* is the post-stimulus time (PST), *t*_0_ is the latency offset (constant delay), *t*_l_ is the latency scaling (cumulative delay), *a*_0_ is the amplitude offset and *a*_l_ is the amplitude scaling – see Figure 1C. Thus for latency, *t*_0_ moves the whole HRF forwards or backwards in time, while *t*_l_ extends the HRF in proportion to the time relative to trial onset. This model has greater flexibility, e.g, in separating amplitude from different types of latency, than the above linear basis sets. It is fit by maximising the correlation between (normalised) template and (normalised) individual FIR. We have previously applied this approach to modelling the effect of age on the latency of evoked MEG responses in the same task and participants (Price et al., 2017), and use the same fitting procedure described there (gradient ascent on Pearson correlation). For similar approaches to nonlinear HRF fitting, see Kruggel & Yves Von Cramon (1999).^5^

#### HDM3 model

This nonlinear model is a biophysical, generative model based on differential equations that can be integrated to produce HRFs, as a function of biologically relevant parameters (e.g, vasodilatory time constants), rather than parameters that simply describe the shape of the HRF (as in NLF4 above). We used a well-established hemodynamic model (Friston et al., 2000; Stephan et al., 2007; Havlicek et al., 2015) with some updates to the parameters. The form of the model is explained in detail in Appendix 5 of Zeidman et al. (2019), and it is shown schematically in Figure 1D. To briefly reprise, there are four hidden states: vasoactive signal *s* regional cerebral blood flow *f_in_*, venous volume *v* and deoxyhemoglobin *q*. Their dynamics are governed by the following equations:

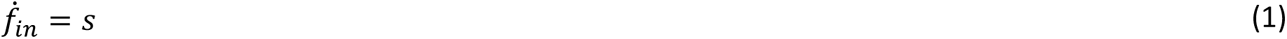

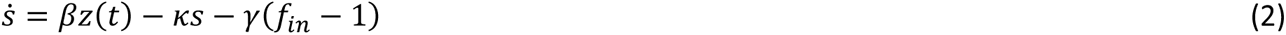

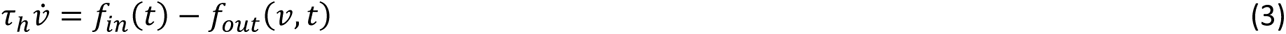

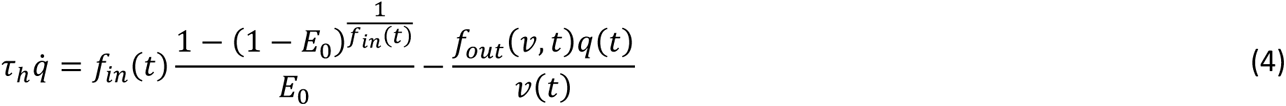

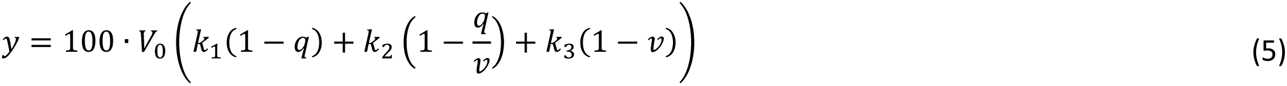

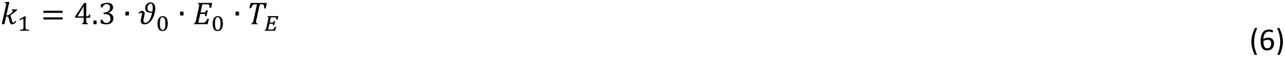

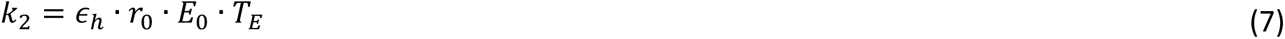

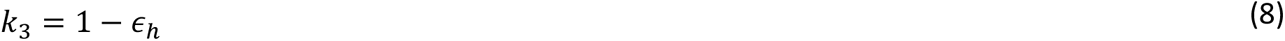

Equations 1-2 model the vasoactive signal *s*, which induces blood flow *f* in in response to neural activity *z*(*t*) ∈ {0,1} with neural efficacy parameter *β*. This part of the model subsumes a variety of neurovascular coupling mechanisms, such as nitric oxide signalling. The decay of the vasoactive signal is governed by rate parameter *κ*, while rate parameter *γ* provides autoregulatory feedback^6^. Together these equations form a coupled oscillator.

Equations 3-4 model the cerebral blood volume *v* relative to the blood volume at steady state, as well as the proportion of deoxyhemoglobin relative to rest, *q*. These equations correspond to the balloon model of Buxton et al. (1998), where parameter 1/*τ*_h_ is the rate of blood flow through the vessel, and parameter *E*_0_ is the resting oxygen extraction fraction. The blood outflow from the vascular compartment is *f_out_* (*v*, *t*) = *v*(*t*)^l/a^, in which Grubb’s exponent *α* relates cerebral blood flow (CBF) and cerebral blood volume (CBV).

Finally, Equations 5-8 model the generation of the fMRI BOLD signal (Obata et al., 2004, Stephan et al., 2007). Parameter *V*_0_ is the resting blood volume fraction, *ϑ*_0_ is the frequency offset at the outer surface of magnetised vessel, *r*_0_ is the slope of intravascular relaxation rate against O_2_, *T_E_* is the echo time and *ϵ_h_* is the ratio of intra-vascular to extra-vascular components of the gradient echo signal.

In the present HDM3 model, we allowed three parameters to be estimated from the data: 1) neural efficacy parameter, *β* (as a proxy for neural activity), 2) rate of decay of vasoactive signal, *κ* and 3) transit rate of blood flow, 1/*τ_h_*. The remaining parameters were fixed based on empirical priors (see Supplementary Material).

To fit the HDM, we used https://github.com/pzeidman/HDM-toolbox, which requires the SPM software package (https://www.fil.ion.ucl.ac.uk/spm/software). The model was fit using the standard Bayesian model fitting scheme in the SPM software, called Variational Laplace (Friston et al., 2007). For participant *i* with timeseries *Y_i_*, this returns a multivariate normal probability density over the parameters, *P*(*θ*_i_** |*Y*_i_**)∼*N*(*μ_i_*, Σ*_i_*). The estimated values of the parameters *μ_i_* were those that maximised the log-evidence for the model, as approximated by the free energy *F_i_* ≈ ln *P*(*Y_i_*).

While this scheme provides parameter estimates for individual participants, *μ_i_*, it also yields the estimated covariance of the parameters Σ*_i_*. To convey both *μ_i_* and Σ*_i_* to the group level, we used the Parametric Empirical Bayes (PEB) framework in SPM (Friston et al., 2016, Zeidman et al., 2019). This is a Bayesian hierarchical linear regression model applied to the estimated parameters of all participants’ models (see Supplementary Material for more details). We then tested the evidence for the presence versus absence of each covariate (mean and effect of age) on each hemodynamic parameter using Bayesian Model Reduction (BMR, Friston et al., 2018), which iteratively prunes mixtures of group-level parameters from the PEB model, where doing so does not reduce the free energy.

### Temporal resolution

In terms of temporal resolution (which determines the PST resolution in Figure 4 and latency resolution in Figure 6), the choice of the FIR bin width of 1s was deemed as a reasonable trade-off between bin-width and number of samples per bin (and this sub-TR sampling is only possible because of the jitter described above; Josephs & Henson, 1999). The Can3 basis functions were simulated at the resolution of 0.062s (TR/32), as stated in the “Single-participant (1st-level) models” section above. Even though the NLF model is fit to the FIR, and so has the same 1s resolution, the latency scaling is estimated via interpolation, so can have higher resolution. Finally, the HDM model was simulated at its default resolution of 0.375s (a higher resolution of 0.062s did not affect results).

### Nonlinearities as a function of SOA

Nonlinearities are known to exist for SOAs below around 10s. A key feature of the HDM3 model is that, as a non-linear model of hemodynamics, it explicitly captures these nonlinearities, which can then be visualised using a mathematical device called a Volterra expansion (Friston et al., 2000). This expresses the relationship between the output of the system (the HRF) and the input of the system (the stimulus) as the weighted sum of *Volterra kernels*. A first-order Volterra kernel is the modelled HRF in response to a brief stimulus, and a second-order Volterra kernel is the nonlinear change in the HRF as a function of time elapsed since the previous stimulus. (Crucially, the second order kernels can only be estimated in the context of an experimental design that samples nonlinearities, through the use of variable SOAs, as here.) These kernels are shown and discussed in Supplementary Figure S8.

Of the total variance in the BOLD timeseries explained by the two Volterra kernels, the proportion explained by the second-order kernel ranged from 4.5% to 29% across ROIs (i.e, the first-order kernel explaining the remaining 95.5% to 71%). Thus nonlinearities as a function of SOA were relatively small, though not insignificant, which needs to be kept in mind when interpreting the results of the other three models that assume linearity. Nonetheless, the proportion of this second-order variance that related to age (linear and/or quadratic) only ranged from 1% to 12% across ROIs, suggesting that this nonlinearity is unlikely to dramatically affect age-related conclusions drawn from the linear models.

## Results

### Whole-brain analyses

Separate analyses were performed to identify BOLD responses time-locked to stimulus presentation in sensory cortices, and BOLD responses time-locked to button presses in motor cortices. ^7^ For each analysis, the parameter estimates for each basis function (time bin) from the FIR32 GLM were entered into a second-level group model across participants to estimate the mean, linear and quadratic effect of age for each basis function.

An F-test across the mean values from the stimulus-locked analysis showed many voxels that responded (relative to interstimulus baseline), the most significant of which were in bilateral auditory and visual cortices (Figure 2A).^8^ Left (contralateral) motor cortex showed a significant mean effect in the response-locked analysis. All three regions showed a linear effect of age, with additional linear effects seen in right (ipsilateral) motor cortex (Figure 2B), as well as supplementary motor cortex (not analysed further). No voxels showed strong quadratic effects of age.

The stimulus-locked, mean FIR estimates from the peak of the left auditory cortex cluster showed a typical BOLD impulse response, peaking around 4-5s and with a more sustained undershoot (Figure 2C, top). Interestingly, the estimates for the effect of age on each time bin (Figure 2C, bottom) showed a profile that resembled a temporal derivative of the mean response, such that the BOLD response became more delayed with age (which can be visualised by adding the bottom plot in Figure 2C to the top plot). Similar results were found for the visual and left motor clusters (not shown, but see later).^9^

The response-locked mean FIR for the rMC (Figure 2D, top), however, was not typical, and it was the effect of age (Figure 2D, bottom) that appeared more like a typical HRF. This pattern can be explained when plotting the HRF by age in the next section, where it becomes apparent that the young showed a negative BOLD response that disappeared with age (see also Mayhew et al., 2022; Tak et al., 2021).

The clusters showing linear age effects were used to define four functional ROIs: lAC, bVC, lMC and rMC (see Methods). FIR models were then re-fit to the first temporal component of a singular value decomposition of each voxel’s BOLD timeseries for each ROI. The first 16s of these ROI FIR responses are shown for each participant as heatmaps in the top row of Figure 3, and averaged within age tertiles as plots in the top row of Figure 4.

**Figure 3.**
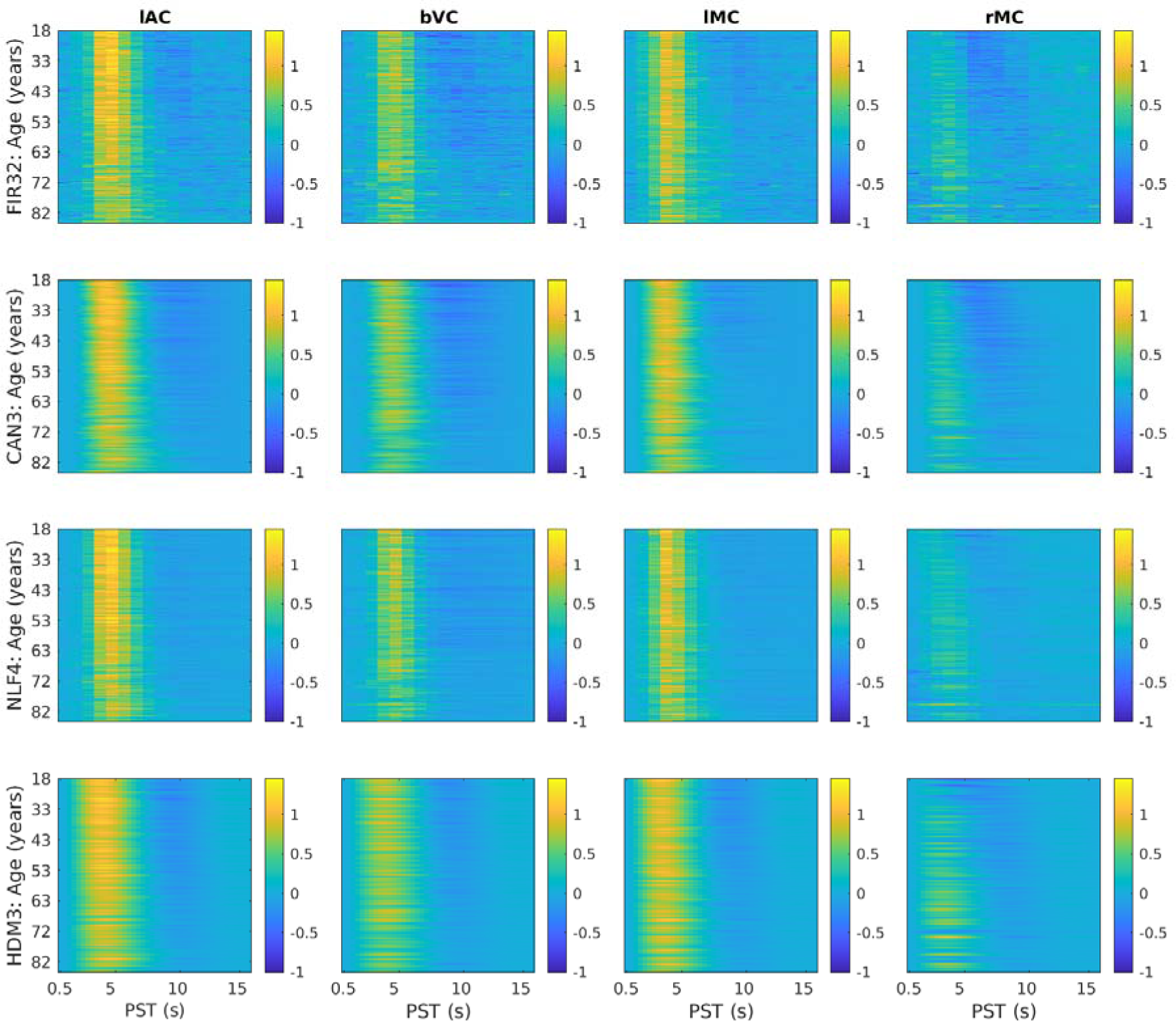
HRFs shown as heatmaps for each ROI (columns) and model (rows). The y-axis represents each participant, sorted by age, while the x-axis represents post-stimulus time (PST), truncated at 16s for easier visualisation. The fits were smoothed across participants with a 5-participant running average. lAC = left auditory cortex, bVC = bilateral visual cortex, lMC = left (contralateral) motor cortex, rMC = right (ipsilateral) motor cortex. The lAC and bVC data are from the stimulus-locked model; lMC and rMC are from the response-locked model.

**Figure 4.**
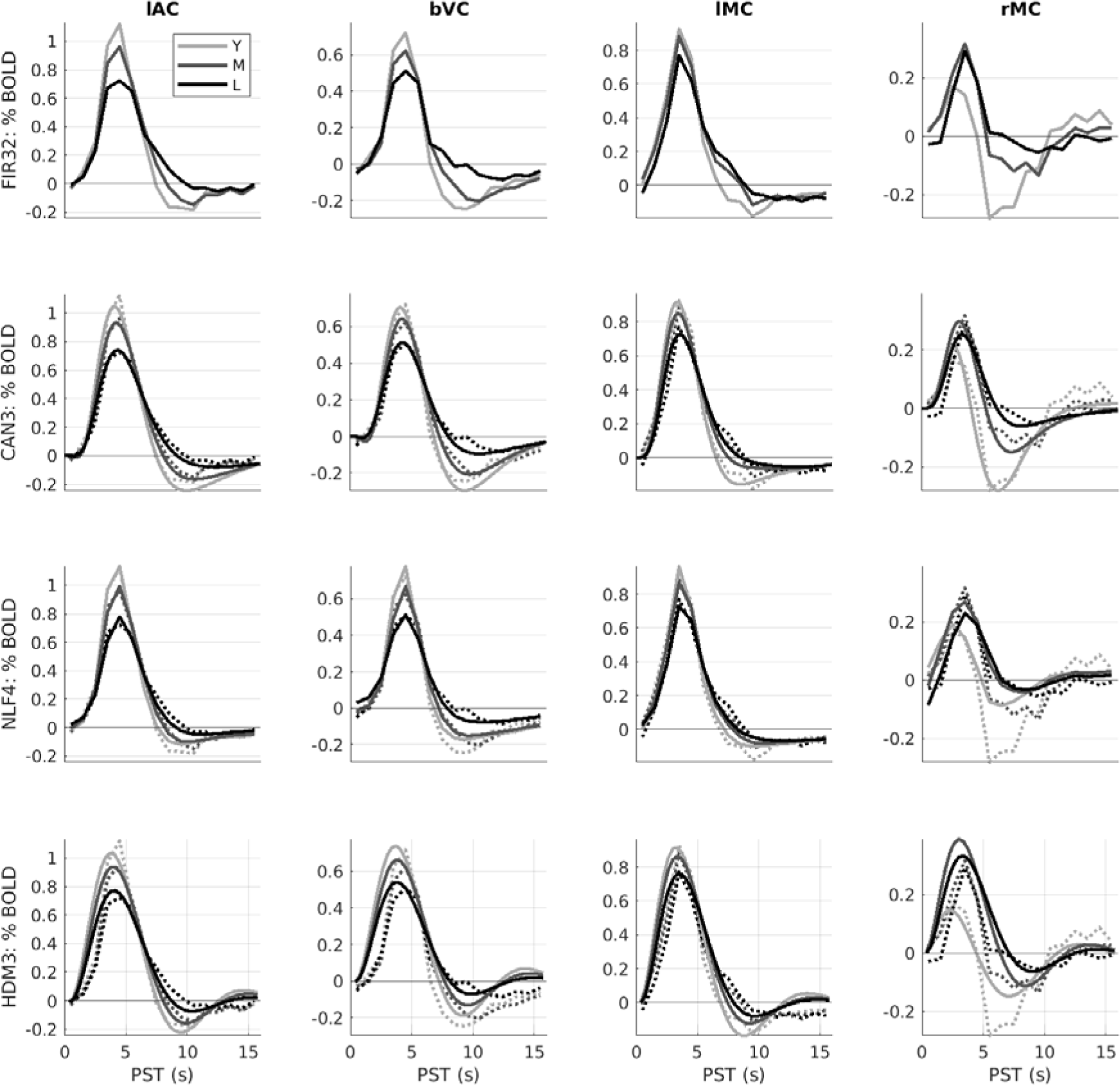
Average HRFs within age tertiles (18-44, 44-66 and 66-88 years) for each ROI (columns) and model (rows). Y = young; M = mid-life; L = late-life. The dotted lines in rows 2-4 are the (replotted) FIR estimates (i.e. same as top row) for reference. Note y-axis scale different across ROIs but matched across models. See Figure 3 legend for more details.

The effect of age for stimulus-locked responses in lAC and bVC is a reduction in amplitude (of both peak and undershoot), and an increase in dispersion (e.g., delay in centre of mass of the response). Indeed, the undershoot almost vanishes for the oldest participants. For response-locked data, the lMC shows a similar increase in dispersion, but smaller age-related decrease in amplitude. The rMC however shows a quite different response that is almost triphasic, with a smaller peak but larger undershoot in young people (explaining the mean and age effects in Figure 2D). Note also that there is some variability in HRF shape across ROIs (see Supplementary Figure S3).

### Features of FIR32 fits

One could extract a number of features (i.e., descriptive statistics) from the FIR fits to each participant. As an example, we defined the peak amplitude and peak latency from the FIR time bin with the maximal absolute value within the first 16s.

For peak amplitude, Spearman rank correlations showed significant decreases in peak amplitude with age for lAC, bVC and lMC, but a significant increase in peak amplitude for rMC (top row of Figure 5).

**Figure 5.**
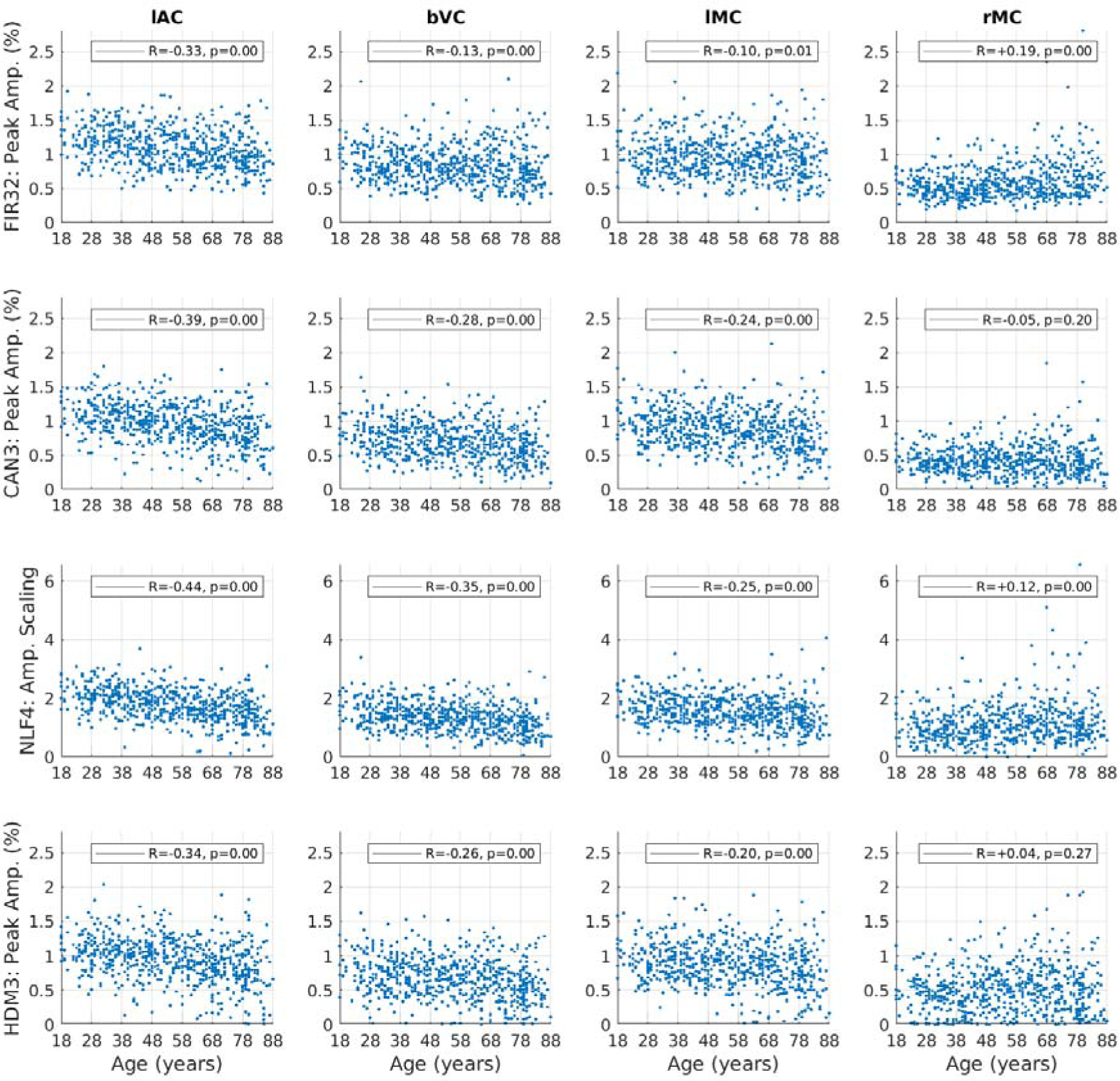
Amplitude estimates for each ROI (columns) and model (rows), plotted for each participant as a function of age. Amplitude is defined as the maximum absolute value of the fit within the first 16s, except for the NLF4 model, where the amplitude scaling parameter is plotted instead (since it is designed to capture amplitude without defining a peak). The R and p-value for Spearman rank correlations with age are shown in legend. See Figure 3 legend for more details.

For peak latency (top row of Figure 6), there is clearly limited resolution (given the FIR bin size) and there are many potential outliers (even when truncated at a maximum of 16s). Nonetheless, Spearman rank correlations showed small but significant effects of age in increasing peak latency for lAC and lMC. Peak latency decreased with age in rMC, though this likely reflects the difficulty of defining the peak for the more complex response shape in this ROI (cf. the latency parameters from NLF model below).

**Figure 6.**
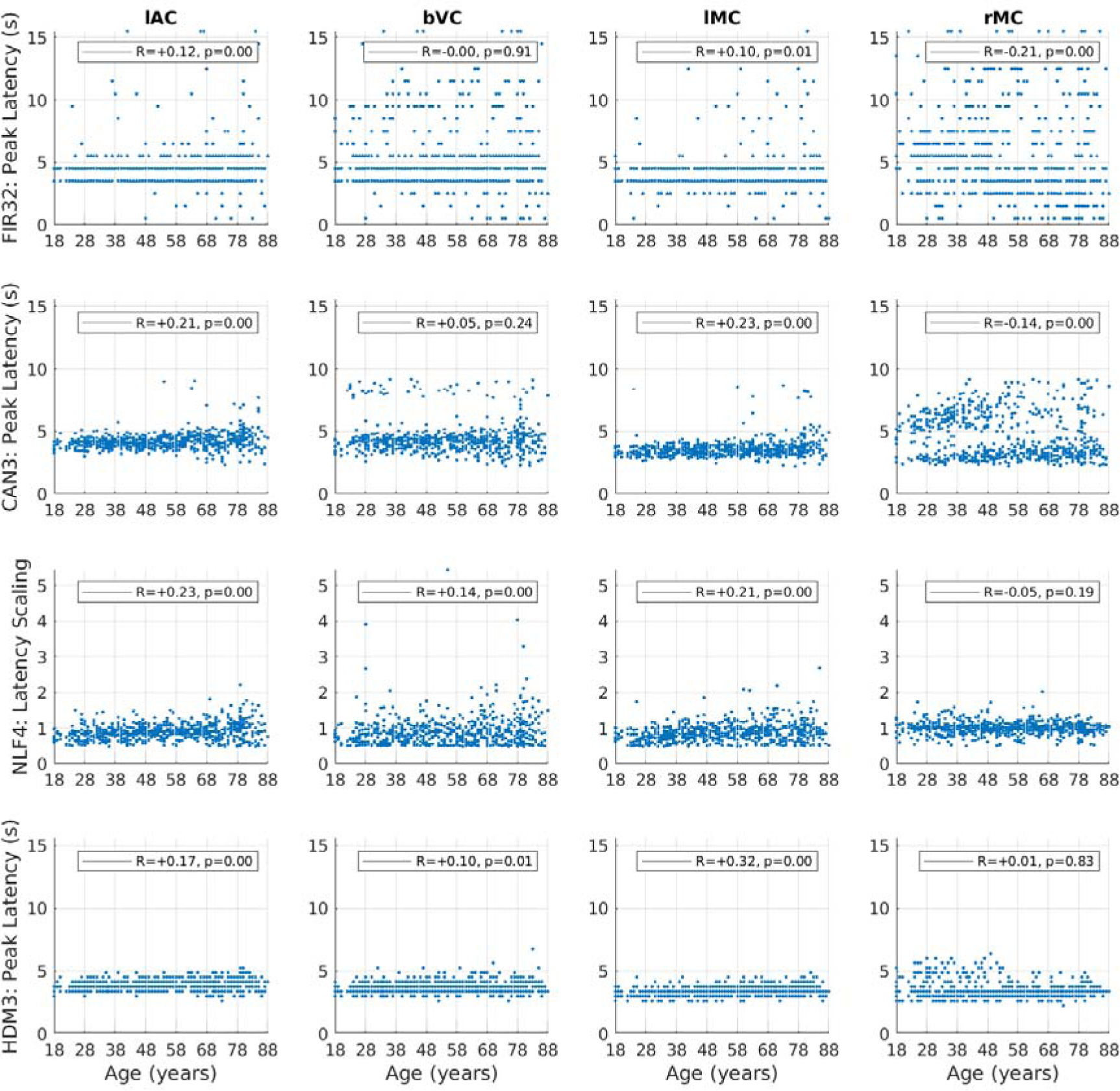
Latency estimates for each ROI (columns) and model (rows), plotted for each participant as a function of age. Latency is determined by the time of the maximum absolute value of the fit (peak) within the first 16s, except for the NLF4 model, where the absolute value of the latency scaling parameter is plotted instead (since it is designed to capture latency without defining a peak). See Figure 5 legend for more details.

We also examined the root-mean squared error (RMSE) of the residuals across the original BOLD timeseries (top row of Supplementary Figure S4a). Note there is some statistical circularity for the FIR model, since the same model was used to define the ROIs in the first place, but its inclusion here at least provides a lower bound on the residuals, albeit biased, with which to compare the other models. Not surprisingly therefore, the FIR32 model had least error in fitting the original timeseries, though also likely because it had the most degrees of freedom (i.e, most flexibility). This error then increased numerically from the Can3 to NLF4 to HDM3 models.

The bottom row of Supplementary Figure S4a shows the RMSE across trial-averaged, post-stimulus time, namely the fit of the Can3, NLF4 and HDM3 models to the FIR fit. In this case, the NLF4 model had least error (except in rMC), followed by the Can3 model and then HDM3.

Note however that a better test of the models than the above RMSE metric is to apply cross-validation, in order to adjust for differences in model flexibility, as is done later. More importantly, Supplementary Figure S4b plots the residuals (from original timeseries) as a function of age. The residual error increased with age for all models and in all ROIs except bVC, as did the variability across participants. This suggests that a subset of older participants have more (non-stimulus-locked) noise remaining in their fMRI data (e.g, residual effects of head motion) and/or demonstrate more trial-to-trial variability in their HRF.

### Can3 fits and HRF features

The second row of Figures 3-4 shows the fits of the Can3 basis set (see Methods), while the second row of Figures 5-6 shows its estimates of peak amplitude and peak latency derived from the reconstructed HRF for each participant. Figure 4 shows that the Can3 basis set is accurate in capturing variation in the BOLD response shape across age tertiles and across ROIs (closely following the FIR estimates in dotted lines). One potential limitation of the Can3 set is the inability to decouple the amplitude of the peak and that of the subsequent undershoot (see, for example, lAC and bVC where the magnitude of the undershoot is slightly over-estimated).^10^

The correlations of peak amplitude and peak latency with age were stronger for the Can3 fits than the FIR fits for lAC, bVC and lMC (cf. first and second rows of Figures 5-6). This suggests that the Can3 basis set is more sensitive to effects of age on features of the HRF shape (as would be expected when the peak amplitude and latency are effectively derived from a weighted sum of many time bins, rather than a single time bin in case of the FIR). For rMC however, Can3 no longer showed a significant increase in peak amplitude with age, and showed a smaller negative correlation for peak latency (cf. the NLF4 model considered next). Indeed, there is clearly a bimodal distribution of peak latencies from the Can3 model, reflecting whether the peak or undershoot has greater absolute displacement from zero.

### NLF4 fits and HRF features

The third row of Figures 3-4 shows the fits of the NLF4 model (see Methods), while the third row of Figures 5-6 shows its estimates of amplitude and latency. This model does not appear to fit all ages as well as the Can3 model, particularly effects of age on the undershoot, and particularly in the rMC. The latter is understandable from the fact that the average across participants (or first singular temporal component) for the rMC may not correspond to the typical “template” HRF required by this approach.

Note that the purpose of the NLF4 model is to estimate amplitude and latency directly, rather than derive them post hoc from the peak of the fitted responses, so the third row in Figure 5 reflects the NLF4 amplitude scaling parameter and the third row in Figure 6 reflects the NLF4 latency scaling parameter. Indeed, this model distinguishes two types of amplitude and latency – an offset (shift) and a scaling (stretch) – see Methods. The advantage of the NLF4 model can be seen in the generally stronger correlations of amplitude and latency with age. For lAC and bVC for example, the amplitude scaling parameter shows a more negative effect of age than does the peak amplitude estimated from fits of the FIR32 or Can3 basis sets, while the latency scaling parameter shows a more positive effect of age. This is because the NLF4 model explicitly separates the amplitude from the latency. For rMC, the NLF4 model recovers a significant positive effect of age on amplitude, but no longer produces a significant negative effect of age on latency, most likely because it eschews the need to define the peak of a more complex evoked response shape (cf. issue of peak and undershoot for Can3 model above).

### HDM3 fits and its parameters

Whereas the NLF4 model is designed to fit features of the BOLD response shape (specifically amplitude and latency), the final model considered here is a biologically-plausible generative model of how that response is caused. While the full HDM potentially has multiple free parameters (see Supplementary Material), we focus on an HDM3 version, in which we fixed several parameters to their prior expected value, leaving three free parameters: neural efficacy, neurovascular decay rate and hemodynamic transit rate.

The bottom row of Figures 3-4 shows the fits of the HDM3 model, while the bottom row of Figures 5-6 shows estimates of amplitude and latency from the HDM3 fit. The HDM3 model does reasonably well for all ROIs, though appears to peak earlier than the FIR model, and struggles to capture age effects in rMC.

The three free parameters of the HDM3 model are shown as a function of age in Figure 7. Neural efficacy (top row) showed a small negative effect of age in lAC and bVC (but not lMC), but a positive effect of age in rMC. Neurovascular decay rate (second row) showed strong positive effects of age in all ROIs. The hemodynamic transit rate (bottom row) showed strong negative effects of age in lAC, bVC and lMC, but not rMC. Indeed, in rMC, there were many participants for whom the default transit rate of 1.02 was sufficient (likewise for the default neurovascular decay rate of 0.64).

**Figure 7.**
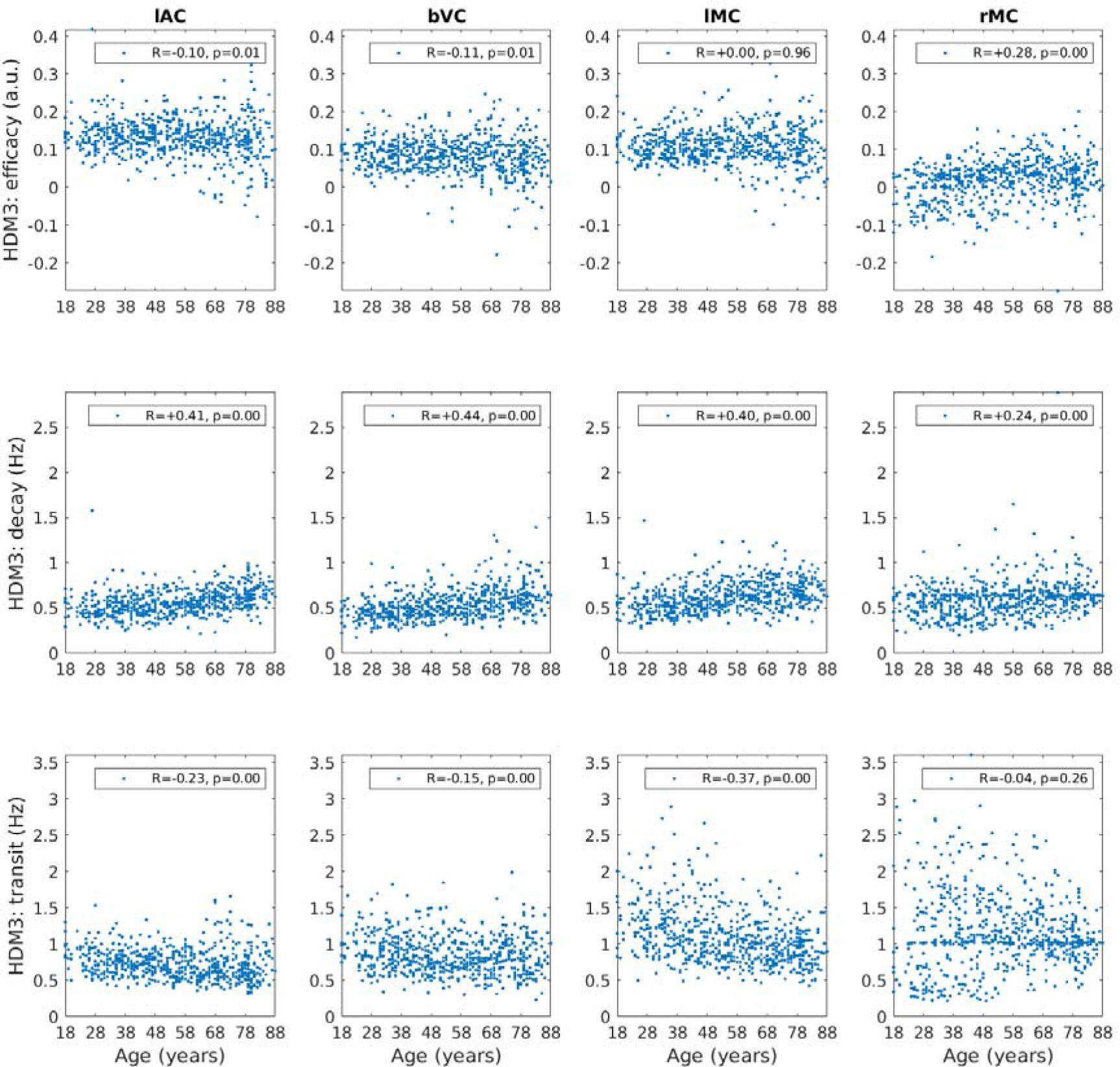
HDM3 parameters (rows) for each ROI (column) plotted for each participant as a function of age. The R and p-value for Spearman rank correlations with age are shown. Parameter estimates for decay and transit rates are after transforming the posterior expected values from their log-values back into original units of Hz (see Supplementary Material).

Furthermore, we can incorporate the HDM results into a hierarchical linear model, in order to regularise the parameter estimates across participants, using SPM’s PEB framework (see Methods). More specifically, we fit a group-level model that included a constant term (average parameter value across participants) and a linear age effect (slope of parameter change with age). We then used BMR to prune parameters that are not needed in order to maximise the model evidence for this group-level model. This analysis takes into account the posterior covariance between parameters, effectively dropping parameters whose effects can be accommodated by another parameter (while maintaining similar model evidence).

The results of PEB-BMR on the linear effect of age on each parameter are shown in Figure 8 (results for the mean across participants are shown in Supplementary Figure S5). After PEB-BMR, the posterior expectation for the neural efficacy parameter remained close to its prior (0) in lAC, bVC and lMC, i.e., there was no need for age to moderate neural efficacy. Only in rMC was there a need to increase neural efficacy with age. By contrast, neurovascular decay rate showed an increase with age across all ROIs, while hemodynamic transit rate showed a decrease with age in lAC, bVC and lMC.

**Figure 8.**
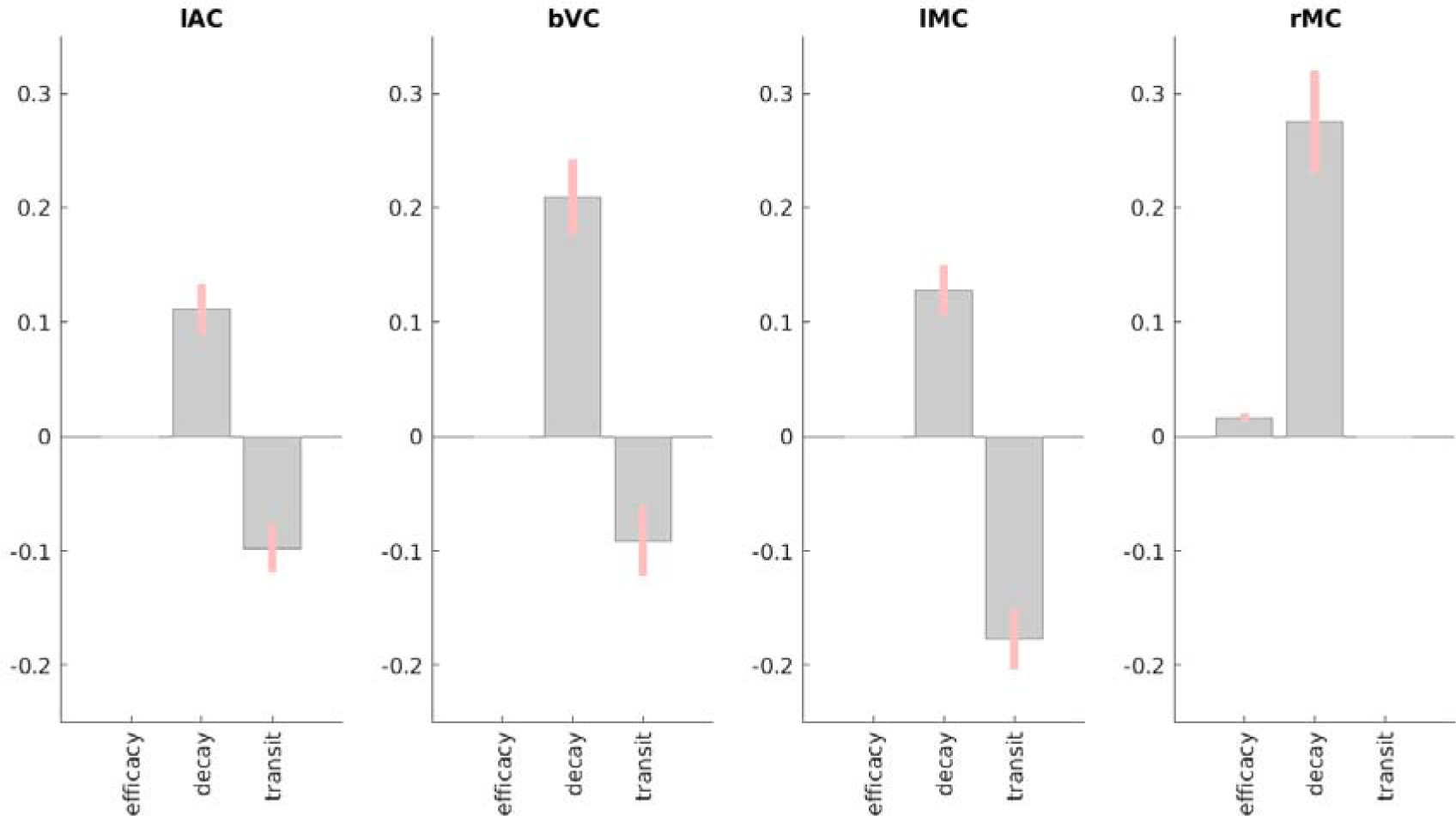
Results of PEB BMR on HDM parameters for each ROI for the linear effect of age across participants. Grey bars show posterior expectation with 90% credible interval in pink; missing bars are parameters that BMR has removed as unnecessary (in terms of maximising evidence for PEB model). For decay and transit parameters, units are log deviations.

Note that while these Bayesian results were generally consistent with the Spearman correlations of each parameter with age in Figure 7, at least for the decay and transit parameters, the PEB-BMR results question the significant negative Spearman correlations of neural efficacy with age in lAC and bVC. This is because the PEB-BMR approach takes into account the posterior covariance among parameters, which was ignored by the correlation analyses above, which were done independently on each parameter. The presence of correlation between the posterior estimates of the neural efficacy parameter estimates and those of the other two “vascular” parameters means that not all are needed to simultaneously capture the effects of age on the BOLD response.

### Cross-validated Prediction of Age

We tested how well each of the four models could predict participants’ chronological age based on their parameter estimates, combined across all four ROIs, using multiple linear regression and leave-one-out cross-validation.^11^ The top panel of Figure 9 shows that all models did a reasonable job of prediction, with the Can3 model explaining the most (44%) variance in age, and the NLF4 model explaining the least (25%).

**Figure 9.**
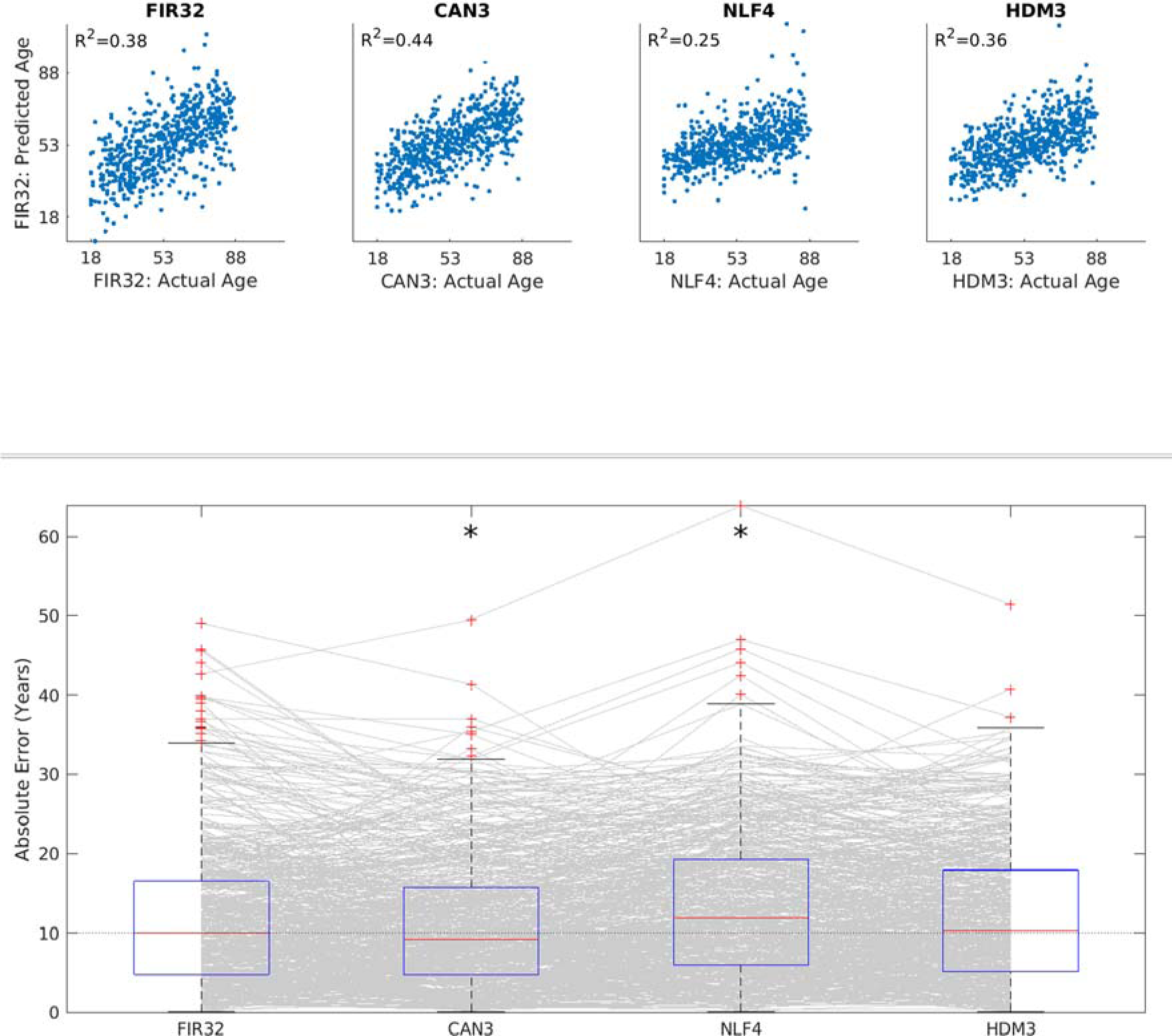
Results of cross-validated prediction of age for each model, using all parameters across all ROIs. Top panel shows the Pearson correlation between actual and predicted age. Bottom panel shows boxplots of absolute error, where each grey line corresponds to one participant. An asterisk means that a two-tailed sign-test revealed significantly different error than from the FIR32 model. The average across ROIs of the median error was 9.98 years (FIR32), 9.20 years (Can3), 11.90 years (NLF4) and 10.33 years (HDM3), respectively.

The bottom panel of Figure 9 shows the absolute error in those predictions. Sign tests showed that the Can3 model did significantly better (smaller median error) than the FIR32 model, despite the fact that the FIR32 model had smaller RMSE when fitting individual timeseries data (see Supplementary Figures S4a and S4b). This is likely because the much greater number of parameters in the FIR32 model results in over-fitting (i.e, capturing some noise in individual fits), such that their estimates do not generalise well to data from other individuals. The NLF4 model was significantly worse than the FIR32 model, suggesting that its parameters were failing to capture some important aspects of ageing. The HDM3 model did not differ significantly from the FIR32 model, which is reassuring for this model, though it did not perform as well as the Can3 model.

For comparison, Supplementary Figure S6 shows the age prediction error for the Can3 model, when fitting the canonical HRF only, or adding its temporal derivative, or adding both its temporal and its dispersion derivatives. This suggests that all three basis functions are needed.

Supplementary Figure S7 shows the age prediction error for four versions of the HDM model (the HDM3 used here, plus versions with 4, 5 and 6 free parameters). The HDM3 model did better than the more complex models with more parameters (indeed, performed significantly better than the HDM4 model). This is why we focused on the HDM3 model here.

### External validation of HDM3 parameters

Finally, we sought external validation of the HDM3 parameters. In previous work (King et al., 2023), we estimated three latent vascular factors (LVFs) from six potential measures of vascular health in the CamCAN dataset. These factors corresponded roughly to total blood pressure (LVF1), pulse pressure (LVF2) and heart rate variability (LVF3), shown to be associated to cerebrovascular health measures in the CamCAN cohort (Fuhrmann et al., 2019; Tsvetanov et al., 2021). N=625 of the present participants had valid data for these LVFs.

A priori, the LVF1 and LVF2 factors that related to blood pressure would seem most likely to relate to the hemodynamic transit rate parameter of the HDM model. Nonetheless, to test for any possible relationship between the three vascular factors and three HDM3 parameters, we performed nine mediation analyses (using the Matlab toolbox available here: https://github.com/canlab/MediationToolbox; Shrout & Bolger, 2002), testing whether each LVF mediated the effect of age on each HDM3 parameter.^12^ The HDM3 parameters were taken before PEB-BMR (see above), and averaged across the four ROIs for each participant, in order to reduce multiple comparisons and potentially obtain a more accurate estimate for each participant, assuming that the true vascular parameters are similar across brain regions within the same person.

The only mediation test that survived Bonferroni correction for nine tests was that of the first LVF (LVF1) on the HDM3 transit parameter. In this model, age was positively related to LVF1 (total blood pressure; a-path=0.34, T=8.97), and negatively related to mean transit rate (c-path without LVF1 present=-0.27, T=-6.90), both as expected. Somewhat surprisingly, LVF1 was positively related to transit rate (b-path=0.15, T=3.64), such that higher total blood pressure was associated with faster transit, whereas one might expect age-related chronic vascular alterations, such as atherosclerosis, to lead both higher BP and lower CBF (slower transit times). In any case, the relationship between age and transit rate became more negative with the inclusion of the LVF1 mediator (c’-path=-0.32), producing the significant mediation effect (ab-path=0.05, T=3.36, p<.001). While interpretation of this mediation may be complex (see Discussion), the fact that the only analysis to reveal a relationship between age, vascular health and HDM parameters involved the expected transit rate HDM parameter and one of the two expected LVFs related to blood pressure, supports the validity of the above age-related changes on HDM parameters.

We also performed mediation analyses relating the HDM parameters to an independent measure of neural activity in the same participants performing the same task in MEG (see Price et al., 2017, for details). We took the mean evoked responses estimated from source-reconstruction of planar gradiometer data from N=586 of the present participants, and summarised these responses in terms of their energy (sum-of-squares of values from 0 - 400ms post stimulus- or response-onset, after mean-correction for a pre-onset baseline period of 100ms).

A priori, we would expect this MEG measure of neural activity to mediate the effect of age on the neural efficacy parameter of the HDM3 model, but not the other two vascular parameters. In this case, we ran mediation analyses separately for each ROI, since the HDM neural efficacy parameter showed different effects of age in the rMC compared to other ROIs. Thus, we ran all twelve possible mediation analyses (three HDM3 parameters and four ROIs). None of these analyses showed a significant mediation effect, even at p<.05 uncorrected (and even in rMC). Thus we could not independently validate the HDM neural parameter with these MEG data, though we note that the energy of first 400ms of the evoked response is unlikely to capture all neural activity detectable by fMRI (see Discussion).

## Discussion

We compared the ability of four models to capture age-related changes in the BOLD response to brief audiovisual stimuli and key presses, using a large (N=645) and representative dataset containing individuals uniformly distributed across the adult lifespan, and an fMRI paradigm optimised to estimate the shape of the HRF.

Whole-brain analysis using the first model – the “FIR32” model with 32 temporal basis functions capturing the HRF every 2s – revealed strong effects of age in many brain regions, particularly bilateral visual, auditory and motor cortices. The predominant effects of increased age were to reduce the amplitude of the BOLD response and to shift its centre of mass later in time.

Focusing on four of these ROIs, we compared the FIR fit to a more parsimonious linear model using a canonical HRF and two of its partial derivatives (“Can3” model), as well as two nonlinear models. In terms of capturing features of the HRF such as peak amplitude and peak latency, the Can3 model showed stronger effects of age than the FIR32 model. However, a nonlinear model in which 4 parameters were fit to the FIR estimates (“NLF4” model, comprising zero- and first-order expansions of amplitude and post-stimulus time) showed even stronger effects of age in most ROIs, suggesting better de-coupling of amplitude and latency.

However, it is unclear whether such HRF features are informative with respect to the underlying neural activity, which is normally what fMRI is used to infer. Indeed, the mapping from neural activity to BOLD response is nonlinear, depending on vascular and hemodynamic parameters that are also likely to change with age. Only the fourth model, based on prior work on biophysical, generative hemodynamic modelling (Buxton et al., 1998; Friston et al., 2000), attempted to estimate such neural and vascular parameters separately. When allowing three of these parameters to vary (neural efficacy, neurovascular decay rate and hemodynamic transit rate), this “HDM3” model suggested that the majority of age-effects on the BOLD response arise from vascular/hemodynamic differences. Indeed, only one ROI – right motor cortex (rMC) – showed an effect of age on HDM3’s neural parameter, and this actually reflected increased neural activity in this ipsilateral motor region with age (see below for further discussion about this particular region).

We compared all four models in terms of the accuracy with which the age of a new participant could be predicted from parameters fit to the remaining participants, i.e., their out-of-sample cross-validated performance. Given that the models are typically fit using different statistical frameworks (ordinary least squares, gradient ascent and Variational Bayes), the cross-validated performance was useful as a common basis for comparing all the models’ sensitivity to ageing. Comparison of the average (absolute) prediction error for 1, 2 and 3 basis functions in the Can3 model (Supplementary Figure S6), and of the four versions of the HDM model with 3-6 parameters (Supplementary Figure S7), confirmed the value of this approach. When applied to the four main models considered here, age prediction was better for the Can3 model than FIR32 model, suggesting that the greater flexibility of the latter resulted in over-fitting. Age prediction for the NLF4 model, on the other hand, was worse than the FIR32 model, suggesting that it cannot capture all types of age-related variance. The HDM3 model was comparable to the FIR32 model, suggesting that it is doing a reasonable job, in addition to providing more physiologically-meaningful parameters.

Finally, we sought external validation of the HDM parameters using independent data about vascular health (abstracted from blood pressure, BMI and ECG measures) and neural activity (from an analysis of evoked MEG responses by the same participants in the same task). While our MEG measure did not provide evidence of mediating the effect of age on the HDM3 neural parameter, one of our independent vascular factors did show evidence of mediating the effect of age on the HDM3 hemodynamic transit time, which provides some validation for the HDM3 model.

### Studying Ageing with fMRI

If one just wants to detect (e.g, localise in the brain) the effects of age on the BOLD response, rather than interpret those effects in terms of neuronal versus vascular influences, then the present results demonstrate that SPM’s canonical basis set, Can3, is best. It outperformed the FIR32 model in its cross-validated ability to predict age, and will offer greater statistical power by virtue of using fewer degrees of freedom.^13^

However, if one wishes to interpret age effects on the BOLD response in terms of neural activity, then such linear basis functions are not sufficient. For this, a biophysically-plausible generative model is needed. Although its performance was worse than the Can3 model, cross-validated age-prediction performance of the HDM3 model was comparable to the FIR32 model. Importantly, if the HDM3 model correctly distinguishes neural and hemodynamic contributions to the BOLD response, then the present results have important potential implications for previous fMRI studies that have interpreted age-related differences in the BOLD response in terms of differences in neural activity (at least in sensorimotor tasks like the present one). This is because the present HDM3 results suggest that most age effects on the BOLD response arise from vascular factors instead. This is consistent with recent work combining BOLD fMRI with glucose PET (Stiernman et al., 2023), which showed that effects of age on the BOLD response (during a working memory task) were not accompanied by effects of age on glucose metabolism. Indeed, we found that age need not affect neural efficacy in early visual, auditory and contralateral motor regions, at least when using a Bayesian approach to prune away parameters whose effects are close to zero or well captured by the other parameters. This is consistent with a previous study (Tsvetanov et al., 2015) that adjusted activations in the present task by empirical estimates of vascular reactivity from an independent resting-state fMRI dataset, called Resting-State Fluctuation Amplitudes (RSFA). That study also found dramatically attenuated effects of age on visual, auditory and contralateral motor BOLD responses, once they were scaled by RSFA. Interestingly, that study also found that the BOLD increases with age in rMC remained after RSFA scaling, again consistent with the present HDM3 results in suggesting a neural origin for age effects in ipsilateral motor responses.

The positive effect of age on the vasodilatory decay parameter of the HDM3 model in all ROIs suggests that vasodilatory signals decay faster in older people. This seems plausible, and may reflect changes in the efficacy of neurovascular coupling. Increasing this decay rate reduces the peak response and attenuates the post-peak undershoot (Supplementary Figure S1). Meanwhile, the negative effect of age on the hemodynamic transit rate parameter in most ROIs (except rMC) suggests that blood flow is reduced (i.e., longer transit times) in older people. This also seems plausible; cerebral blood flow decreases 0.3-0.5% per year in healthy ageing (Graff et al., 2023). Decreasing this hemodynamic transit rate delays the HRF (Supplementary Figure S1).

The HDM3 finding that age increases the neurovascular decay rate is also consistent with a previous study that used DCM to examine resting-state connectivity in the CamCAN dataset (Tsvetanov et al., 2016), in which there was a significant positive effect in three of the four resting-state networks. That study also found the same negative effect of age on the hemodynamic transit rate, though it was only significant in one resting-state network. Nonetheless, that study used a different neural model (based on a 1/f neural power spectrum in the resting-state, rather than the brief burst of neural activity in response to a stimulus assumed here) and a different formalisation of the HDM (in which the ratio of intra-vascular to extra-vascular components of the signal, *ϵ_h_*, was also estimated; see Supplementary Material). Moreover, it examined different (higher-order) brain regions. These converging findings for the effects of age on the BOLD response are reassuring.

Note that we are not claiming that there are absolutely no age-related differences in neural activity in the present sensorimotor task, particularly given that we have previously shown age-related delays in evoked responses measured by MEG on the same participants in the same task (Price et al., 2017). Indeed, we used the same NLF4 model in that study to demonstrate that age increased the latency offset (shift) in visual evoked responses and the latency scaling (stretch) in auditory evoked responses. However, these latency differences were small (approximately 20ms offset and 20% scaling), and so unlikely to have much effect on the HRF, given that the HRF integrates over several seconds of neural activity. Thus we are claiming that age may not affect the type of neural activity *that can be detected by* fMRI, at least within the present ROIs and type of task. In other words, we are claiming that our HDM3 results demonstrate that neurovascular and vascular changes are *sufficient* to explain age effects on the BOLD response in the ROIs and task considered here. By extension, the onus is on researchers who wish to use BOLD fMRI to infer about effects of age on neural activity to first rule out age-related vascular effects.

### Linear Basis Sets and HRF Features

While the Can3 basis set showed best cross-validated prediction of age, the partial derivatives of the canonical HRF are important to capture this variability (Supplementary Figure S6). Thus, studies of ageing that use only a single canonical HRF are likely to miss age-related differences in the HRF. Such studies need to ensure that parameter estimates for the derivatives are also taken forward to any group-level (2^nd^-level) analyses (or else combined to estimate a shape-independent effect size, e.g., Calhoun et al., 2004; Cignetti et al., 2016). The additional flexibility provided by the partial derivatives is also important to capture variability across brain regions (Supplementary Figure S3).

Supplementary Figure S3 also shows that the canonical HRF used in SPM is more dispersed than the BOLD responses estimated here. One reason for this could be that the canonical HRF was derived from studies in which the data were not slice-time corrected (Friston et al., 1998, 2000), so there was likely to be a delay (around TR/2 on average, typically corresponding to 1-1.5s), relative to stimulus onset, in the activated brain regions. Here we were careful to synchronise the GLM with the reference slice, ensuring no such delay. Another reason is that only a small number of participants were used in those studies, who may not have been representative, whereas the HRFs in Supplementary Figure 3 are averaged across a larger range of adults, spanning the whole adult lifespan. In case it is helpful, we have added a “revised” canonical HRF (and its partial derivatives) to the github page associated with this paper, in which the parameters of its two gamma functions are estimated by fitting the average FIR across participants and across the lAC, bVC and lMC ROIs. However, while this canonical form may be more appropriate for the average, healthy adult aged between 18 and 88 than SPM’s current one, the main point of the present study is to show how age changes the HRF, so ideally researchers would use age-appropriate HRFs for their analyses. By also providing the FIR fit for all current participants on the github page, researchers can define their own canonical HRF matched to the ages of their own sample (at least for the 4 ROIs considered here; or indeed download the raw data and estimate an FIR fit for any set of voxels).

Beyond localisation, it is unclear whether there are effects of age on the shape of the HRF that can be directly interpreted (at least without a biophysical model). There may be information about the latency of neural activity in the latency of BOLD responses, but this seems most likely for latency differences between two or more conditions (e.g. types of stimuli), found within the same brain region and same individual tested within the same session. In such situations, one can reasonably assume that hemodynamic variables are constant, so any differences are neural in origin. In this case, temporally-extended basis sets like the Can3 can better estimate latencies than individual FIR time bins, which are more prone to noise, though the present results show that non-linear fitting of models that explicitly parametrise different types of latency (e.g, offset and scaling, like in the NLF4 model) leads to even higher sensitivity. Even so, reverse inferences about the latency of neural activity from latency of the BOLD response are complicated due to convolution (e.g, a difference in peak latency of the HRF can arise from differences in the duration rather than onset of neural activity, Henson et al., 2002), not to mention potential nonlinearities in the neural-BOLD mapping. Most importantly however, for the study of ageing, or other individual/group differences, it seems unlikely that differences between individuals in the latency (or indeed amplitude) of their BOLD responses could ever be uniquely attributed to differences in neural activity, at least in the absence of a validated biophysical model or independent data about hemodynamic differences.

Conversely, it is important to remember that any individual differences in the HRF (e.g. as a function of age) cannot be uniquely attributed to hemodynamic factors, unless one knows that neural activity is invariant across individuals (or can estimate that activity separately, e.g. with a biophysical model). This point was made by Grinband et al. (2017) in the context of ageing. We believe that our paradigm minimised factors that could cause age-related differences in neural activity (compared to some previous paradigms) because our trial-based design entailed very brief, simple, meaningless stimuli (<=300ms) that required minimal processing. While older people are generally slower to react in such tasks, we accommodated this delay (for analysis of motor cortices) by locking analysis to the keypress rather than the stimulus, which should allow for age-related variability in execution time at least (though differences could remain in motor force and other kinematic variables). However, our task did involve long and variable SOAs, in order to maximise the ability of fMRI to estimate HRF shape (Josephs & Henson, 1999), so it is possible that age-related differences in sustained attention existed that modulated neural activity (Grinband et al., 2017). But this is exactly why more sophisticated models like the HDM are needed (in the absence of independent measures of neural/hemodynamic factors), to try to separate effects of age on neurodynamics and hemodynamics.

### Caveats with HDM

Though the present HDM3 model offers the potential to separate neural and vascular contributions to the BOLD impulse response, this is subject to several assumptions, as detailed in the Supplementary Material. We fixed several parameters of the full HDM to their prior expected values, whereas in reality they may also have changed with age. The reason for fixing some of them is that the effects of several parameters on the BOLD response are very similar, so they are difficult to disentangle in the absence of independent data (e.g, about blood flow or blood volume). This is reflected in high posterior covariances between some parameters when fitting a 6-parameter HDM (Supplementary Figure S2), and the fact that freeing up more than 3 HDM parameters did not significantly improve cross-validated age prediction (Supplementary Figure S7). Thus, interpretation of some of the age effects that we found with the HDM3 model could be wrong, mandating follow-up studies that complement the BOLD response with other measurement modalities.

For example, regarding the neural efficacy parameter, its effects on the scaling of HRF are very similar to those of the venous blood volume fraction (*V*_0_), and *V*_0_ is known to change with age. Nonetheless, since *V*_0_ must be positive, it cannot be solely responsible for the negative BOLD peak we found in rMC. The effects of the neural efficacy parameter can also be similar to those of the resting oxygen extraction fraction *E*_0_ (Supplementary Figure S1), resulting in a high negative correlation in their parameter estimates (Supplementary Figure S2). However, we argue in the Supplementary Material that effects of age on oxygen extraction fraction are unlikely to be detectable in the BOLD signal measured here. The effects of the neural efficacy parameter are also related to vessel stiffness parameter *α* (Grubb et al., 1974), though in this case, they tend to have opposite effects on the HRF amplitude (inducing positive correlation in their parameter estimates). Nonetheless, our application of PEB to the HDM3 model resulted in no effects of age on neural efficacy β in any ROI except rMC, and the latter is unlikely to be explained by parameters such as blood volume fraction, oxygen extraction or vessel stiffness, because the effects of age on these are unlikely to differ dramatically across cortical regions, particularly across contralateral ROIs like lMC and rMC.

Likewise, the effects of age that we did find on neurovascular decay rate and hemodyamic transit rate could also be attributed to effects of age on other HDM parameters that we fixed, particularly the neurovascular feedback parameter, whose effects on the latency of the HRF are similar to those of the hemodynamic transit rate (Supplementary Figure S1). To properly decouple these HDM parameters, one would need independent measurements of blood flow, oxygenation, vasodilatory signal and neural activity, to simultaneously constrain their values, for example using Arterial Spin Labelling (ASL) fMRI for blood flow, or Vascular Space Occupancy (VASO) fMRI for blood volume.

Here we made a preliminary attempt to validate the HDM parameters against other, independent data from CamCAN, using mediation analysis. We did find that the effect of age on the HDM transit rate parameter was significantly mediated by a previous cardiovascular factor related to total blood pressure (systolic plus diastolic). A relationship between hemodynamic transit and blood pressure would appear to make sense. Nonetheless, we note that this mediation took the form of a “suppressor effect”, whereby the negative relationship between age and transit rate became more negative in the presence of the cardiovascular mediator. One interpretation of this is that increases in blood pressure compensate for age-related reductions in transit rate (given the positive relationship between the cardiovascular factor and transit rate). Whatever the interpretation, this highly significant mediation provides some construct validity to the HDM model.

We did not find any relationship between any of the HDM parameters and the amplitude of early, evoked MEG responses from the same participants performing the same task. One might have expected a mediation of age effects on the HDM neural efficacy parameter, but given that the BOLD response integrates over several seconds of neural activity, it is possible that this short-lived evoked amplitude measure is not sufficient. Future work could examine the effects of age on the power spectrum of neural responses (evoked and induced), for example to see if BOLD amplitude relates to a shift in power towards higher frequencies (Kilner et al., 2005).

Thus in summary, we are not claiming that the HDM3 model used here is the best or most accurate model. Rather our aim is to illustrate how biophysical models like the HDM could be used in principle to better interpret effects of age on the BOLD impulse response, e.g. in separating neural and vascular components. Indeed, we expect future studies to continue to improve the HDM as an explanation for the genesis of BOLD fMRI, and to extend the model to integrate complementary imaging modalities. To illustrate, Havlicek et al. (2015, 2017) proposed improvements to the form of the HDM model to better explain transient features of the BOLD response, and extended it to explain functional ASL measurements. Thus while the present work supports the face validity and predictive validity of HDM, further support for its construct validity will require independent measurement of several of its variables, beyond the final BOLD response. For example, EEG/MEG can provide independent measurements of neural activity from the same people doing the same task, while other types of fMRI contrast, which depend on blood flow or blood volume, could be acquired concurrently.

### Other caveats

There are other caveats with the general approach adopted here. Firstly, while our paradigm (specifically the m-sequence distribution of SOAs) was optimised to estimate the shape of the HRF, this is under linear convolution assumptions. Nonlinearities have been shown for SOAs below around 10s (Friston et al, 2000), which we replicated here from the HDM3 model (see Methods). Furthermore, we showed significant effects of age on the second-order Volterra kernel (Supplementary Figure 8). However, the proportion of variance explained by the second-order kernel was relatively small, and the proportion of this variance related to age was even smaller. Thus we think it unlikely that many of the present inferences about age effects were confounded by nonlinearities as a function of SOA. Nonetheless, this remains possible, and there are other types of nonlinearities, such as nonlinearities in the BOLD response as a function of stimulus duration or magnitude, which are typically bigger than nonlinearities as a function of the SOAs (Birn et al., 2001). While stimulus duration was fixed here, it remains possible that the duration of induced neural activity differed with age. We have no way to test this, other than to note that reaction times (RTs) did not differ much with age in the present task, by design, because it was unspeeded.

Secondly, we identified voxels within ROIs that showed a significant effect (at p<.05 uncorrected) of stimulus/response-locked activity versus baseline. The number of such voxels did decrease with age. While the number of voxels does not necessarily bias estimates of the average across voxels, it is possible that the range of cortical areas sampled, and hence HRF shapes, differed for younger versus older people. Future studies could perform more detailed analyses of individual voxels, possibly better functionally-aligned across participants (than with the present anatomical normalisation), whose HRF shapes might vary, for example, in the contribution of draining veins.

### Ipsilateral motor cortex

Finally, the BOLD response in rMC is noteworthy in its difference from the other ROIs: it showed a negative and delayed BOLD response versus baseline in young people, which became more positive with age (see also (Mayhew et al., 2022; Ward et al., 2008; Tak et al., 2021; Knights et al., 2021, or Mattay & Weinberger, 1999, for review). Indeed, the HRFs in rMC were the hardest to fit, particularly for the HDM3 model. There may be several reasons for this. For example, it has been suggested that the motor cortex ipsilateral to the effector is inhibited by the contralateral motor cortex, but that this inhibition decreases with age, explaining the increase in neural efficacy with age disclosed by the HDM3 model. Such activity in inhibitory interneurons may have metabolic consequences (hemodynamics) that differ from those assumed by the HDM for excitatory activity (Vazquez et al., 2018). Indeed, the neural component of the HDM could be augmented by distinguishing excitatory and inhibitory cell populations, as was proposed for DCM (Marreiros et al., 2008). Furthermore, the neural activity in ipsilateral motor cortex in such tasks could be more delayed/dispersed than the brief burst assumed by all models used here. Ipsilateral motor cortex would therefore be an interesting region to explore more closely in terms of its neurodynamics and hemodynamics.

### Summary

Age has strong effects on the form of the HRF, as shown using all four models we tested, and this should be taken into account when analysing fMRI data. The model one would select for a study depends on its aims. Where the priority is regional localisation, our results demonstrate that the three parameter Can3 model works well. It is parsimonious, has similar sensitivity to ageing effects as the other models we compared, and has best predictive validity. However, when interpreting differences between individuals that have been localised this way, such as due to their age, caution should be exercised before interpreting those differences in terms of neural differences (rather than vascular ones). By contrast, where the aim is to ask mechanistic questions about the underlying causes of ageing effects, there is an opportunity to deploy physiological models such as the HDM. As illustrated here, these kinds of models can generate mechanistic hypotheses, which could then be tested with other modalities that can provide direct measures of neural activity, blood perfusion, or blood volume. Thus, with the appropriate model and data, we can go beyond descriptive statistics such as time-to-peak or peak amplitude, and ask mechanistic questions about the genesis of the BOLD response and its dysfunction in ageing.

### Data and Code Availability

The raw functional and structural images and their preprocessed versions are available from https://camcan-archive.mrc-cbu.cam.ac.uk/dataaccess. All Matlab code for extracting data from ROIs and fitting the models above is available in https://github.com/RikHenson/AgeingHRF.

## Author contributions

Designed research: R.N.H. Performed research: R.N.H., W.O. & P.Z. Analysed data: R.N.H., W.O. & P.Z. Writing—original draft: R.N.H. & P.Z. Writing – review & editing: R.N.H., W.O., K.A.T. & P.Z.

## Supporting information

All Sup Mat

## Acknowledgements

We thank the Cam-CAN respondents and their primary care teams in Cambridge for their participation in this study, and colleagues at the MRC Cognition and Brain Sciences Unit MEG and MRI facilities for their assistance. Further information about the Cam-CAN corporate authorship membership can be found at http://www.cam-can.com/publications/Cam-CAN_Corporate_Author.html (list #14). For the purpose of open access, the author has applied a Creative Commons Attribution (CC BY) licence to any Author Accepted Manuscript version arising from this submission.

## *Cam-CAN corporate author

Lorraine K Tyler, Carol Brayne, Edward T Bullmore, Andrew C Calder, Rhodri Cusack, Tim Dalgleish, John Duncan, Richard N Henson, Fiona E Matthews, William D Marslen-Wilson, James B Rowe, Meredith A Shafto; Karen Campbell, Teresa Cheung, Simon Davis, Linda Geerligs, Rogier Kievit, Anna McCarrey, Abdur Mustafa, Darren Price, David Samu, Jason R Taylor, Matthias Treder, Kamen A Tsvetanov, Janna van Belle, Nitin Williams, Daniel Mitchell, Simon Fisher, Else Eising, Ethan Knights; Lauren Bates, Tina Emery, Sharon Erzinçlioglu, Andrew Gadie, Sofia Gerbase, Stanimira Georgieva, Claire Hanley, Beth Parkin, David Troy; Tibor Auer, Marta Correia, Lu Gao, Emma Green, Rafael Henriques; Jodie Allen, Gillian Amery, Liana Amunts, Anne Barcroft, Amanda Castle, Cheryl Dias, Jonathan Dowrick, Melissa Fair, Hayley Fisher, Anna Goulding, Adarsh Grewal, Geoff Hale, Andrew Hilton, Frances Johnson, Patricia Johnston, Thea Kavanagh-Williamson, Magdalena Kwasniewska, Alison McMinn, Kim Norman, Jessica Penrose, Fiona Roby, Diane Rowland, John Sargeant, Maggie Squire, Beth Stevens, Aldabra Stoddart, Cheryl Stone, Tracy Thompson, Ozlem Yazlik, Dan Barnes, Marie Dixon, Jaya Hillman, Joanne Mitchell, Laura Villis.

## Supplementary Material

### Further details about Haemodynamic modelling (HDM)

Equations 1-8 in the main paper summarise the HDM. Here we expand on its estimation, priors and parameter effects.

Within the model, latent or hidden variables are divided into two types: *states,* which change over time, and *parameters*, which are constant. Only the parameters are estimated from the data, using a Bayesian modelling scheme (Variational Laplace), and the parameters determine the evolution of the states over time.

For the main HDM3 model in the paper, we allowed three parameters to be free: one to capture the magnitude of neural activity, β; one to capture neurovascular effects, namely the rate of decay of vasoactive signal, *κ*; and one to capture vascular effects, namely the transit rate of blood flow, 1/*τ_h_*. The remaining parameters were fixed at their default priors, for reasons expanded below.

- An important vascular parameter is vessel stiffness (the windkessel effect), which is characterised by Grubb’s exponent *α* (Grubb et al., 1974). In the range estimated here, increasing *α* primarily decreases the amplitude of the BOLD response (as well as delay its peak somewhat; see Supplementary Figure S1). Thus its primary effect on the HRF is inversely related to that of the neural efficacy parameter, β, resulting in a positive correlation between their parameter estimates (see Supplementary Figure S2; in general, parameters whose effects are negatively correlated tend to have estimates that are positively correlated, and vice versa). Therefore, we decided to fix the value of *α* here at 0.33 in the HDM3 model, consistent with previous studies (Friston et al., 2000) (though allowed it to vary in the HDM4 model below). This value of 0.33 is consistent with previous human and animal studies, in which values for *α* typically range between 0.2 and 0.4 (as reviewed by Leung et al., 2008). Nonetheless, this parameter is likely to change with age. In rats, *α* has been found to drop from around 0.35 in young individuals (4-5 months) to around 0.25 in very old age (40-41 months) (Dubeau et al., 2011). Thus, it should be kept in mind that some effects of age, particular on the neural parameter *β*, could also be accounted for by differences in vessel stiffness. Then again, our finding using the HDM3 model that there are no effects of age on neural efficacy *β* in any ROI except rMC, is unlikely to be explained by *α*, because vessel stiffness is unlikely to differ dramatically between contralateral ROIs like lMC and rMC (see Discussion of main paper).
- The neurovascular parameter *γ* is the rate constant controlling the feedback from blood flow. Together with the decay parameter *κ*, the two produce a damped oscillation of the vasodilatory signal. However the primary effect of decreasing *γ* is to delay the HRF, which is similar to the primary effect of decreasing the haemodynamic transit rate 1/*τ_h_* (Supplementary Figure S1), resulting in a high negative correlation between them (Supplementary Figure S2). For the HDM3 model, we therefore fixed *γ* here based on its prior expected value of 0.41Hz from Friston et al. (2000) (though allowed it to vary in the HDM5 model below). This does mean, however, that changes in neurovascular feedback could also reflect changes in haemodynamic transit rate.
- Resting oxygen extraction fraction *E*_0_ refers to the percentage of the oxygen removed from the blood by tissue during its passage through the capillary network, and is typically assigned a fixed value (40%) in the HDM model (Friston et al., 2000). The effect of *E*_0_ on the magnitude of the HRF is complex: it can produce an initial negative dip with high values, but at least for values below 40%, increases in *E*_0_ tend to increase the amplitude of the peak of the HRF (Supplementary Figure S1). This can explain why the present empirical estimates of *E*_0_ had a high negative correlation with estimates of the neural efficacy parameter, β (Supplementary Figure S2). While evidence suggests that *E*_0_ increases by about 0.1% per year of adult life (Peng et al., 2014; Leenders et al., 1990), simulations of such changes of up to 7% around the prior of 40% (given the 70 year span in the current sample) showed little effect on the HRF. Therefore we decided to keep *E*_0_ fixed at 40% in the HDM3 model (though allowed it to vary in the HDM6 model below).
- Venous blood volume fraction *V*_0_ is the proportion of tissue occupied by venous blood, and is also typically assigned a fixed value (4%) in the HDM model (Friston et al., 2000). For grey matter containing small vessels, *V*_0_ is generally taken to be in the range 1-4% (Buxton et al., 1998; Havlicek et al., 2015; Hua et al., 2019). Leenders et al. (1990) investigated effects of ageing on *V*_0_, and found it decreases by around 0.05% per year of adult life: from 6.48% for 18 year olds to 3.25% for 88 year olds. Because *V*_0_ simply scales the BOLD response (see Eq. 5), it is perfectly correlated with the neural efficacy *β*, i.e., one cannot estimate both *V*_0_ and *β* using fMRI data alone. Here we chose to estimate *β* instead, but it should be kept in mind that any effects of ageing on *β* could be accounted for by *V*_0_ (though again, our finding of a negative BOLD response in rMC cannot be explained soley by *V*_0_, since *V*_0_ is always positive).
- We adjusted the values of parameters *ϵ_h_*, *r*_0_ and *ϑ*_0_ to correspond to the MRI field strength for the CamCAN data, where *B*_0_ = 3*T*. In previous implementations of this model, the ratio of intra-vascular to extra-vascular signal, *ϵ_h_*, was estimated from the data, reflecting uncertainty about its value in the literature (Stephan et al., 2007). Here, we decided to fix its value for stability, as large values of *ϵ_h_* can induce phase transitions that give rise to unrealistic BOLD responses. Following Havlicek et al., 2015, we used the expected value of *ϵ_h_* = 0.44, based on the ranges of T2* relaxation rates (for further detail, see the MATLAB script *Heinzle_epsilon_derivation.m* in the HDM Toolbox, https://github.com/pzeidman/HDM-toolbox). We set the value of *r*_0_ = 110*s*^-1^ and *ϑ*_0_ = 28.265*B*_0_ = 31.27*s*^-1^, following Heinzle et al. (2016), who drew on the results of Uludag et al. (2009).

**Supplementary Figure S1.**
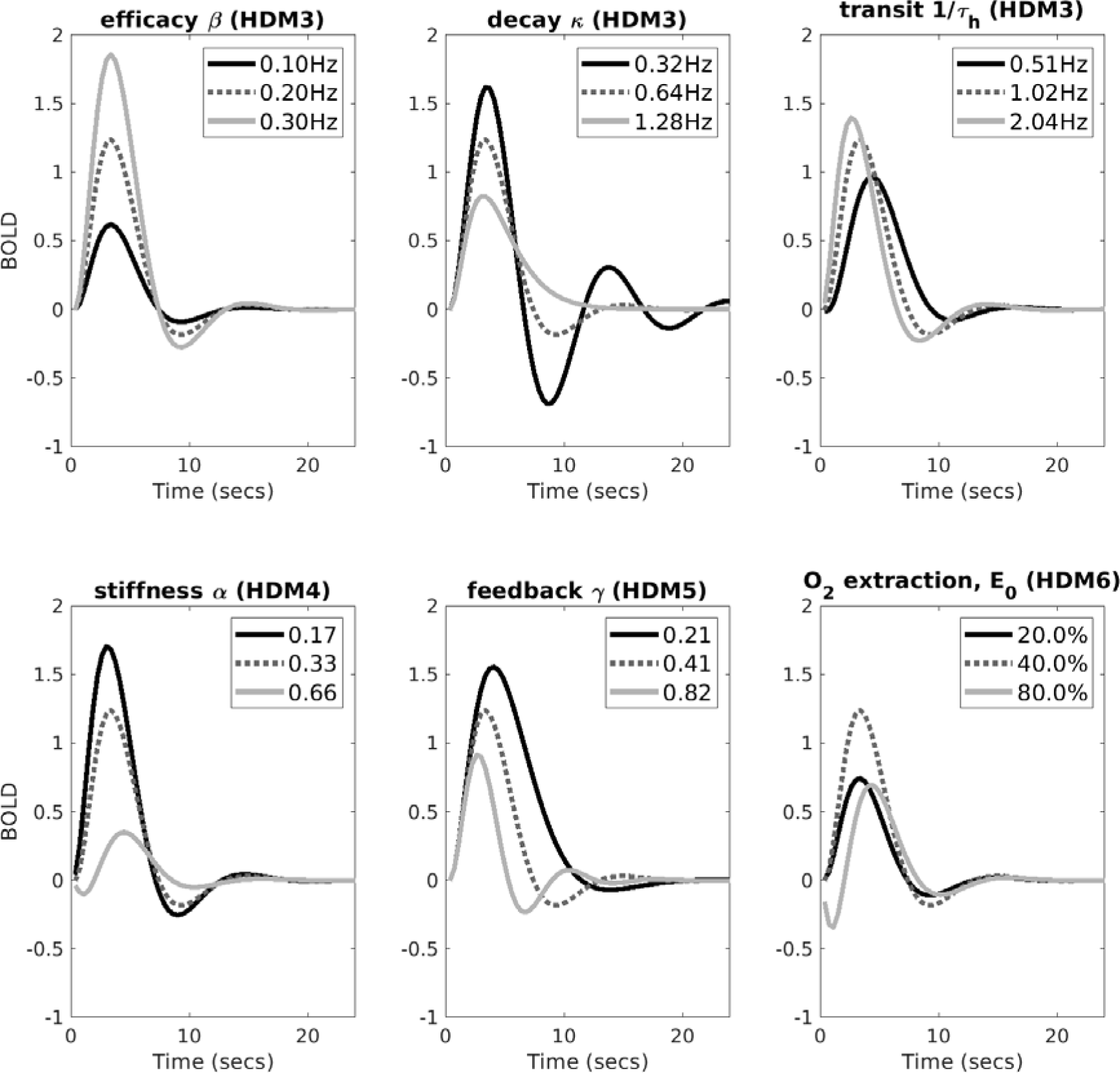
Each plot shows the predicted BOLD response under different values of a parameter, indicated by the title. The central parameter value (producing the dotted line) reflects the prior expectation; the dark solid line and light solid line reflect smaller or larger values respectively (the values of all parameters other than the one varied in a plot were fixed at their prior expectation, except neural efficacy, β, which was set to 0.2).

**Supplementary Figure S2.**
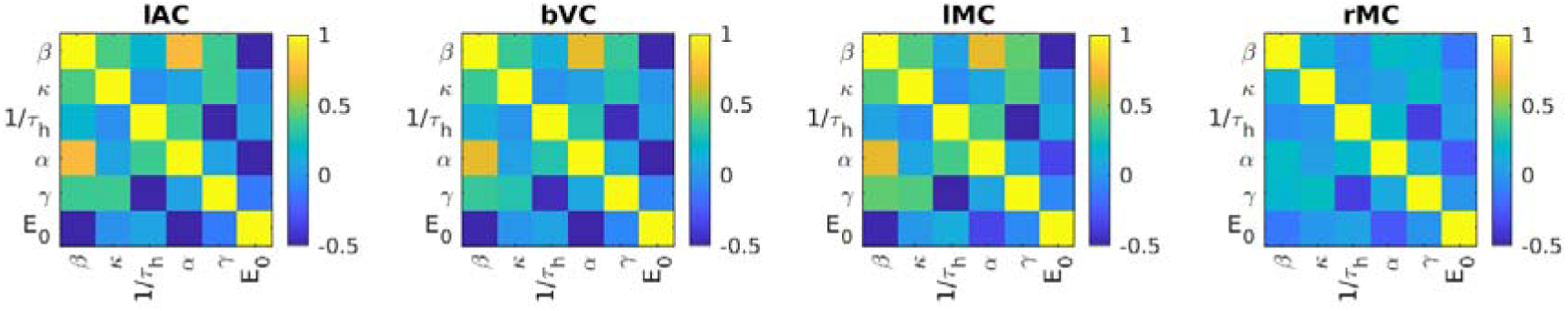
The mean across participants of the posterior correlation between HDM parameters after fitting a 6-parameter HDM model (HDM6) to the data in the paper. Apart from rMC, note that the other ROIs have a large positive correlation between neural efficacy *β* and vascular stiffness *α*, and large negative correlations between neurovascular feedback *γ* and haemodynamic transit rate 1/*τ*_h_, and between oxygen extraction *E*_0_ and *β*.

A summary of the prior expected value and variance for the three free parameters in the main HDM3 model used in the paper are shown in Supplementary Table 1. Note that, in order to enforce positivity constraints on two of these parameters, new parameters *l_K_* and *l_rh_* are introduced, which are log versions of decay rate *κ* and transit rate, 1/*τ_h_*, respectively. The scaling values of 0.64 and 1.02 are taken from (Friston et al., 2000). The neural efficacy parameter *β* is not constrained to be positive so is untransformed.

Additionally, in order to enforce positivity constraints on the states *f*_in_, *v* and *q*, each variable is log-transformed, requiring the equations to be supplemented as follows:

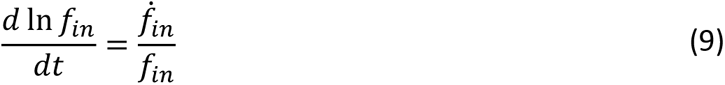

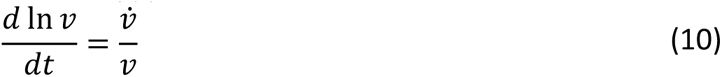

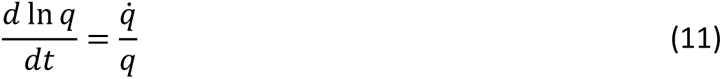

**Supplementary Table 1:**
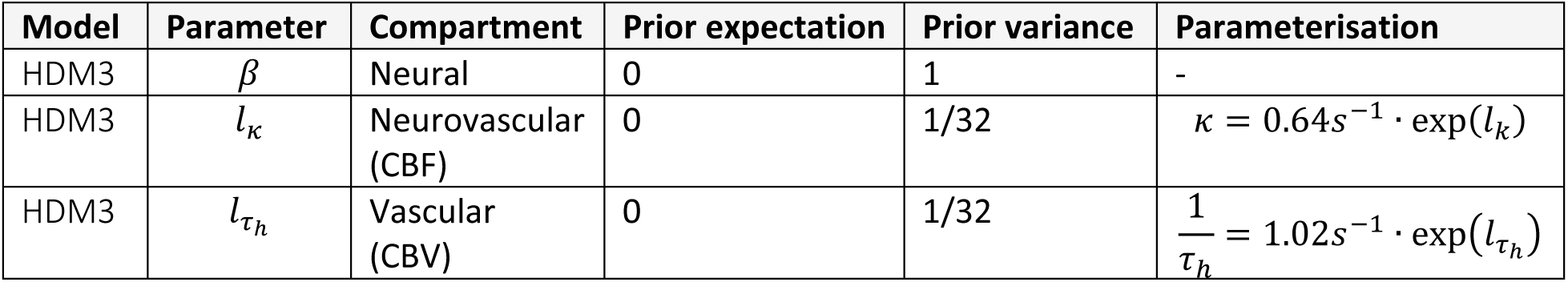
Free parameters of HDM3 model.

The HDM models with 4, 5 and 6 free parameters (used later in Sup Mat) included the values in Supplementary Table 1, plus the one or more of rows of Supplementary Table 2.

**Supplementary Table 2:**
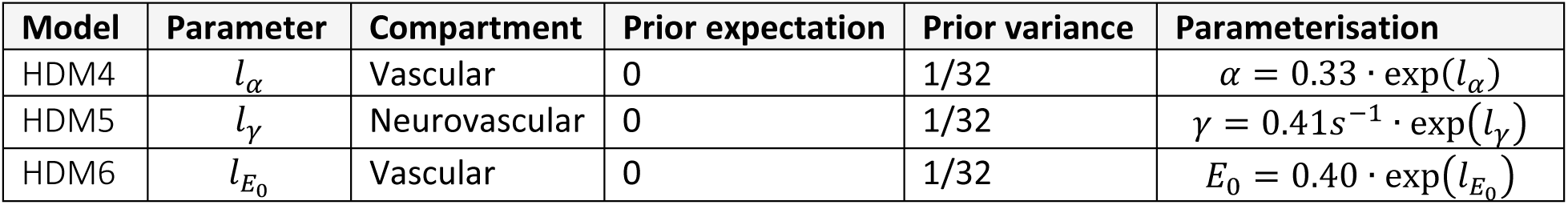
Additional free parameters of HDM4, HDM5 and HDM6 models.

### Group-level model using Parametric Empirical Bayes (PEB)

For the PEB estimation of a group-level model (Zeidman et al., 2019), parameter estimates from all *n* participants are concatenated into a vector of random variables, *θ*^(1)^ = (*θ*_1_, *θ*_2_ … *θ_n_*). Between-participant effects were then modelled using a general linear model:

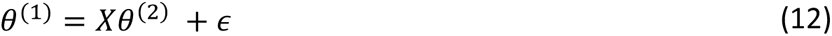

The design matrix *X* included regressors for the effect of each of two covariates (mean over participants and age) on each of *P* = 3 haemodynamic parameters (*β*, *l_K_*, *l_rh_*). This design matrix can be written formally as:

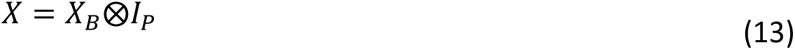

where between-participants design matrix *X_B_* contained two columns: a column of ones (to model the average haemodynamic parameters over participants) and the z-scored age of each participant. This was replicated over the three parameters by taking the Kronecker tensor product ⨂ with the identity matrix of dimension *P*.

Unexplained between-participants variability *ϵ* was modelled using a covariance component model, with a separate I.I.D. precision component for each of the three haemodynamic parameters, allowing each type of parameter to have a separate level of between-participants variability.

The parameters *θ*^(2)^ and free energy *F* of the PEB model were estimated using standard routines in the SPM software. We then tested the evidence for the presence versus absence of each covariate on each haemodynamic parameter using an automated procedure. This was an automatic search that iteratively pruned mixtures of group-level parameters *θ*^(2)^ from the PEB model, where doing so did not reduce the free energy. This was performed using an analytic approach called Bayesian Model Reduction (Friston et al., 2016).

**Supplementary Figure S3.**
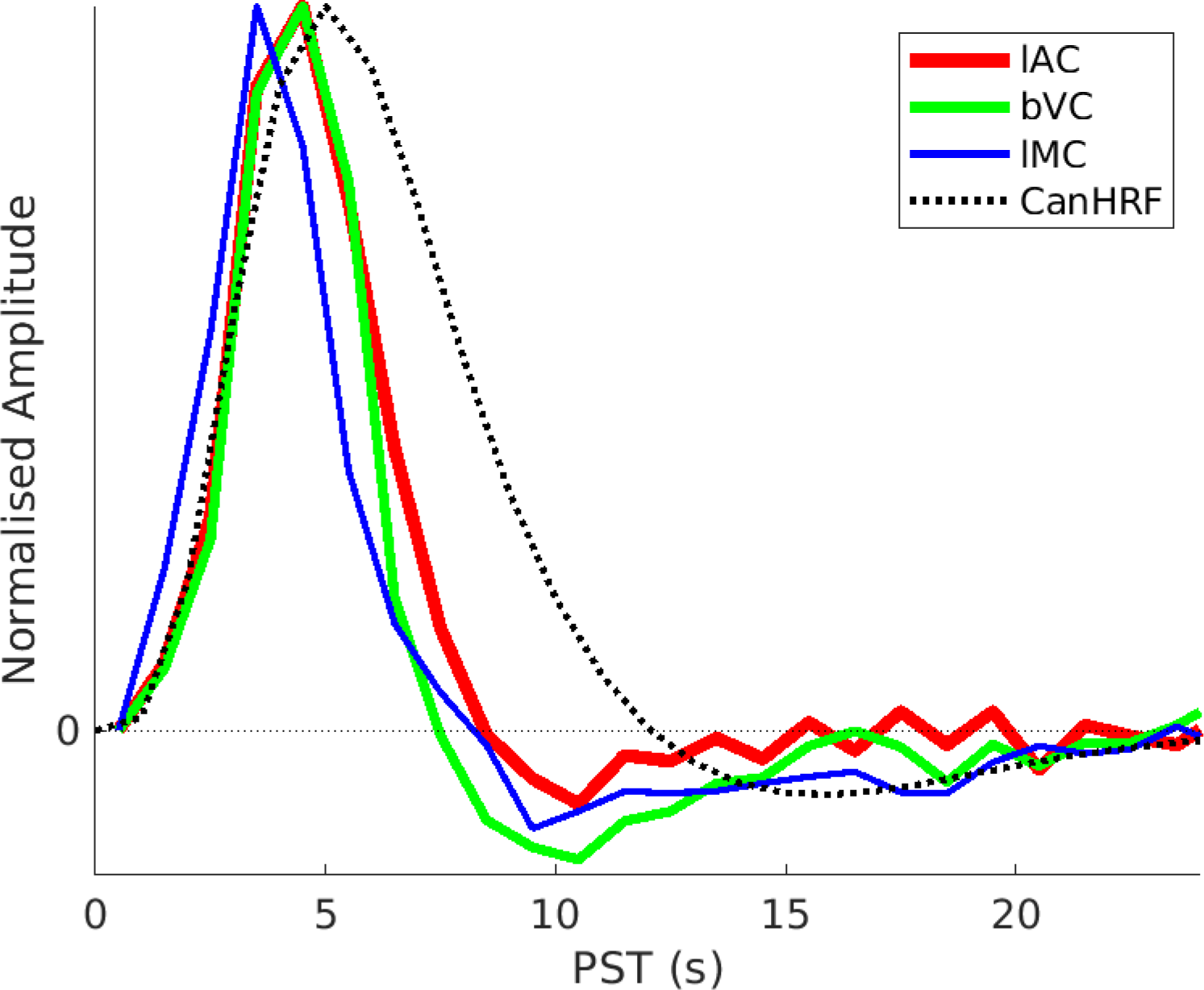
Mean across all participants of the FIR fits for lAC, bVC and lMC (solid lines), along with SPM’s canonical HRF (dotted line). The rMC is not shown because it varied so much with age (see main text). Peak amplitude is matched by scaling by the maximum positive value. The lAC and bVC data are from the stimulus-locked model, while lMC is from the response-locked model. SPM’s canonical HRF is more dispersed than the FIR fits (delayed peak and undershoot). This may not have been noticed before because early studies generating this canonical shape (Friston et al., 2000) did not allow for slice-timing delays. So a delay of TR/2 (typically ∼1s) could partly explain this mismatch. Here we did correct the data for different slice-times and were careful to synchronise the GLM (for FIR32 and Can3 basis sets) with the reference slice used for slice-timing. Grinwald et al. (2017) still observed deviations despite slice-timing, though ones that derivatives should be able to accommodate.

**Supplementary Figure S4a.**
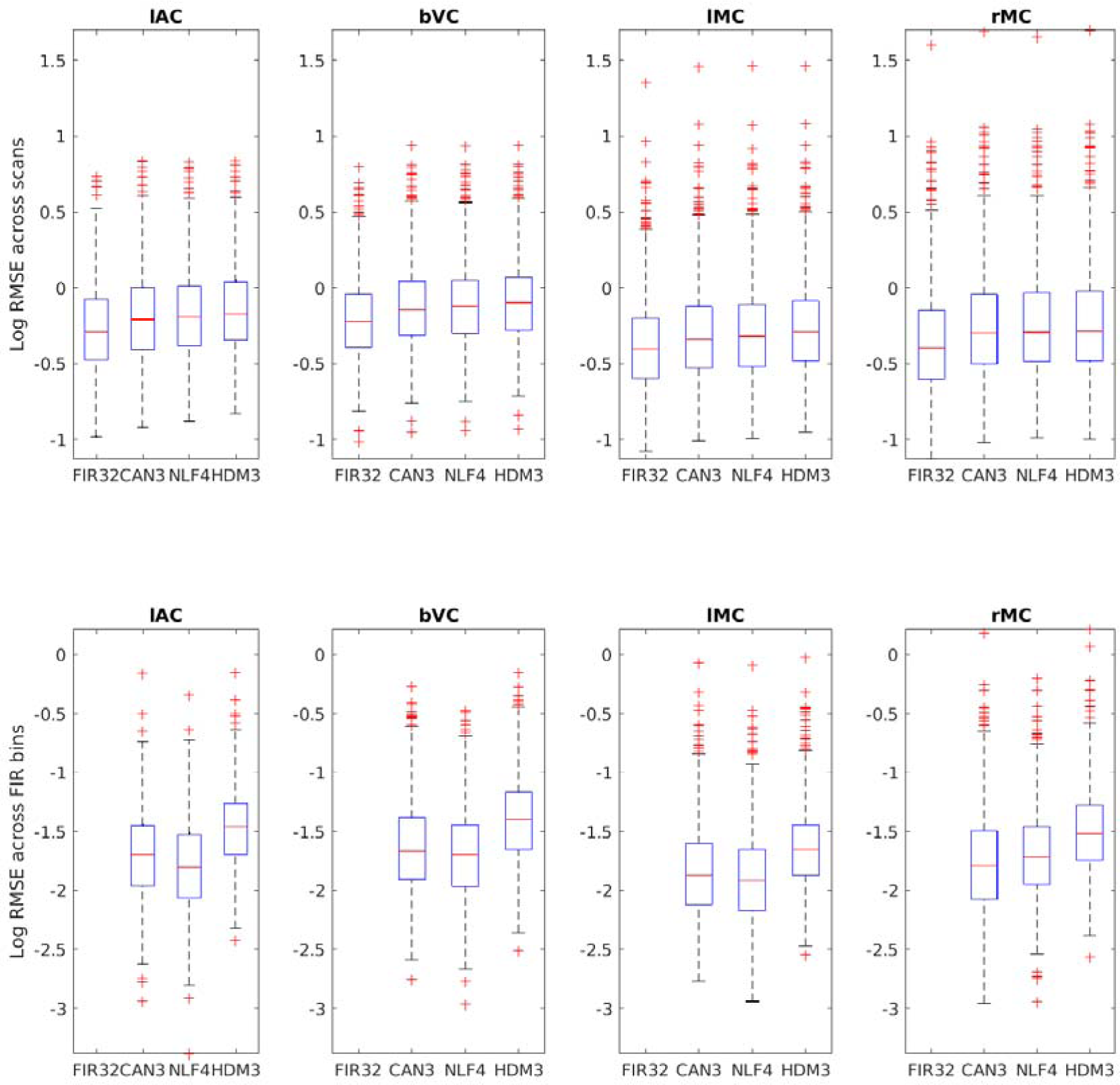
Boxplots (of log) of Root of Mean of Squared Error (RMSE) across scans (top row) or across FIR bins (bottom row) for each ROI and model. Note there is statistical circularity for the FIR model, since the same model was used to define the ROIs in the first place, but its inclusion here at least provides a lower bound, albeit biased, with which to compare the other models. Note also that there can be no RMSE for the FIR32 model in bottom row, and in the top row, the RMSE across scans for the NLF model was calculated by re-inserting the participant- and ROI-specific fitted HRF into each participant’s first-level GLM (hence only 1 effective degree of freedom). The lAC and bVC data are from the stimulus-locked model, while lMC and rMC are from the response-locked model. Note that these measures of model fit ignore differences in model complexity, and so are not as good for generalisation to new data as the cross-validated error in Figure 9.

**Supplementary Figure S4b.**
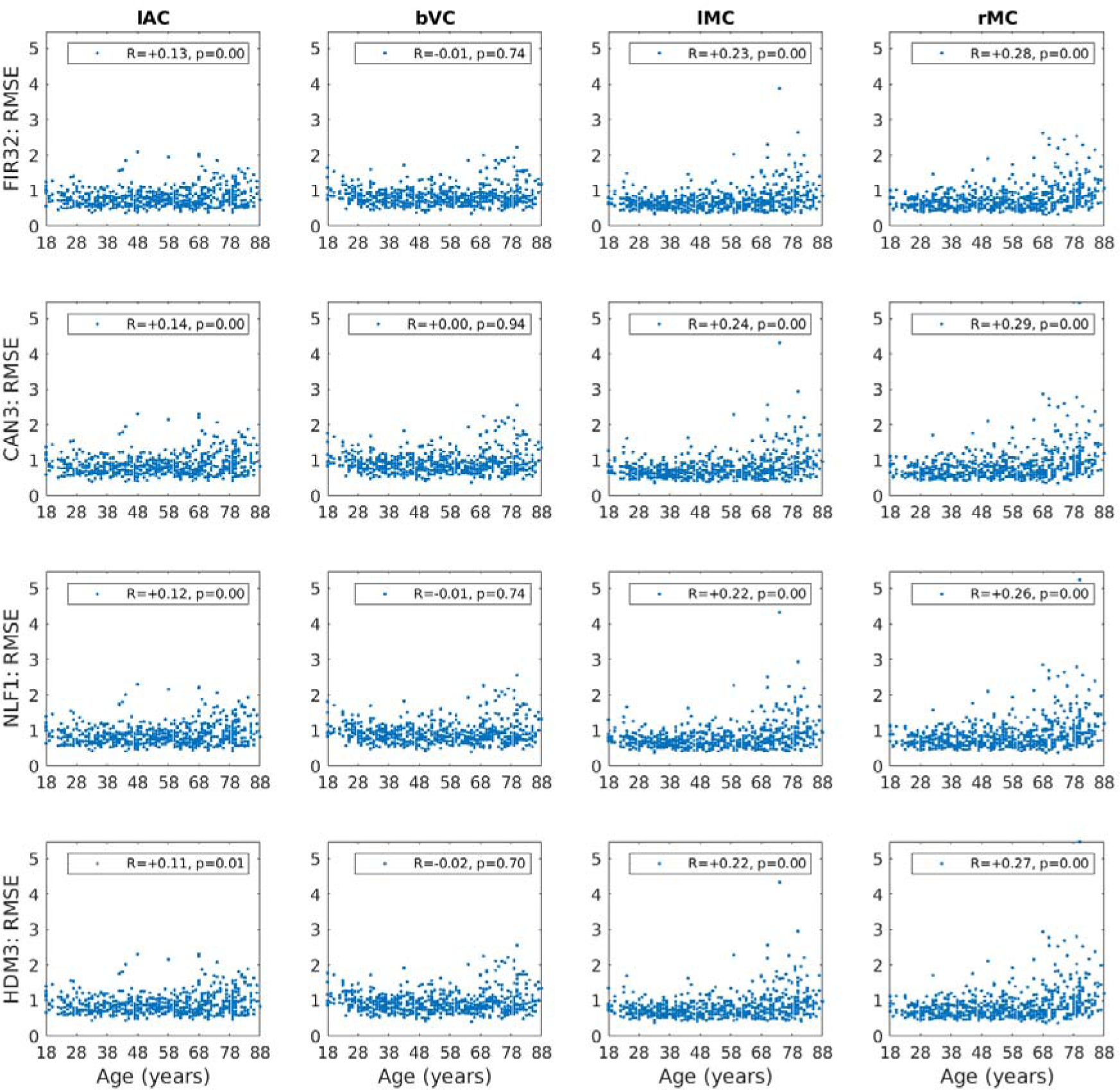
Square-root of Mean of Squared residuals (RMSE) across scans for each model and for each ROI, as a function of participant’s age, together with Spearman correlation R- and p-values. The lAC and bVC data are from the stimulus-locked model, while lMC and rMC are from the response-locked model. Note that the RMSE for the NLF model was calculated by re-inserting the participant- and ROI-specific fitted HRF into each participant’s first-level GLM (hence only 1 effective degree of freedom).

**Supplementary Figure S5.**
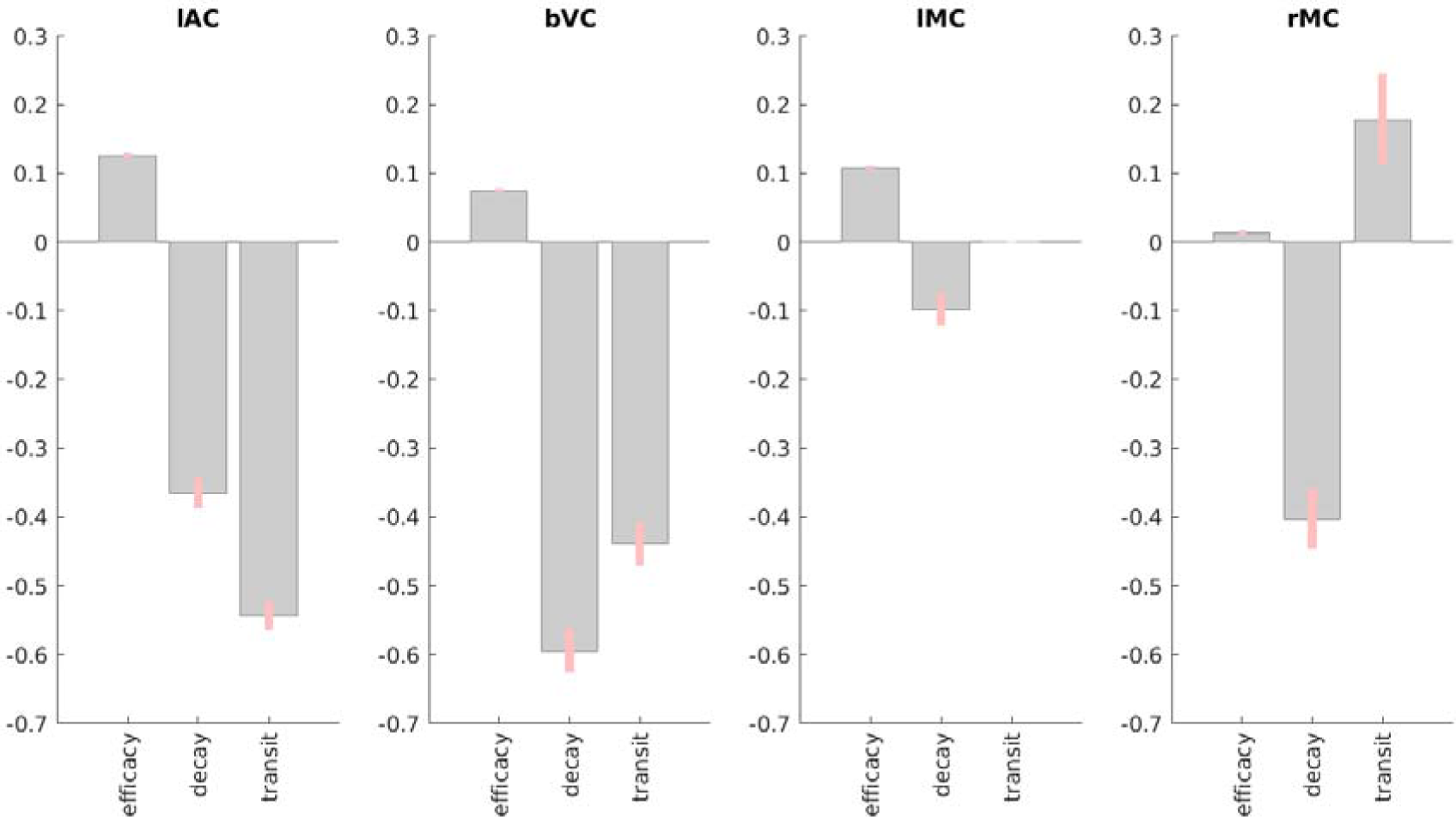
Results of PEB BMR on HDM parameters for each ROI for the mean across participants, in terms of deviation of each parameter from its prior expectation. Grey bars show posterior expectation with 90% credible interval in pink; missing bars are parameters that BMR has removed as unnecessary (in terms of maximising evidence for PEB model). For decay and transit parameters, units are log deviations.

**Supplementary Figure S6.**
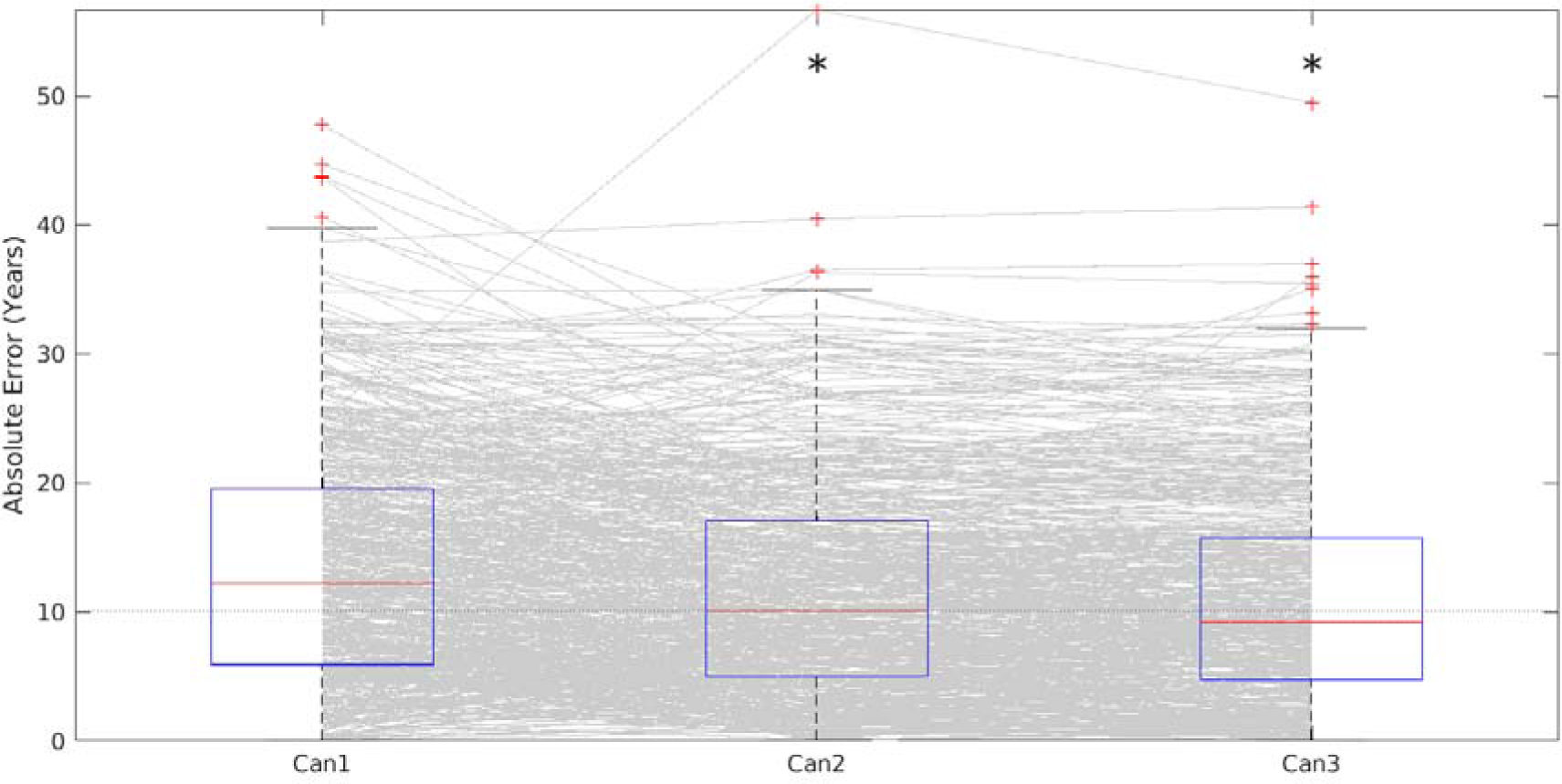
Cross-validated absolute error in predicting age, using Can model parameters combined across all ROIs, as the number of basis functions increases, from canonical HRF only (Can1), to adding its temporal derivative (Can2) and then also its dispersion derivative (Can3). Each grey line corresponds to one participant. Superimposed on these lines are boxplots together with outliers (red crosses). An asterisk means that a sign-test revealed significantly better prediction than the simplest (Can1) model. The median error was 12.2 years (Can1), 10.0 years (Can2) and 9.2 years (Can3).

**Supplementary Figure S7.**
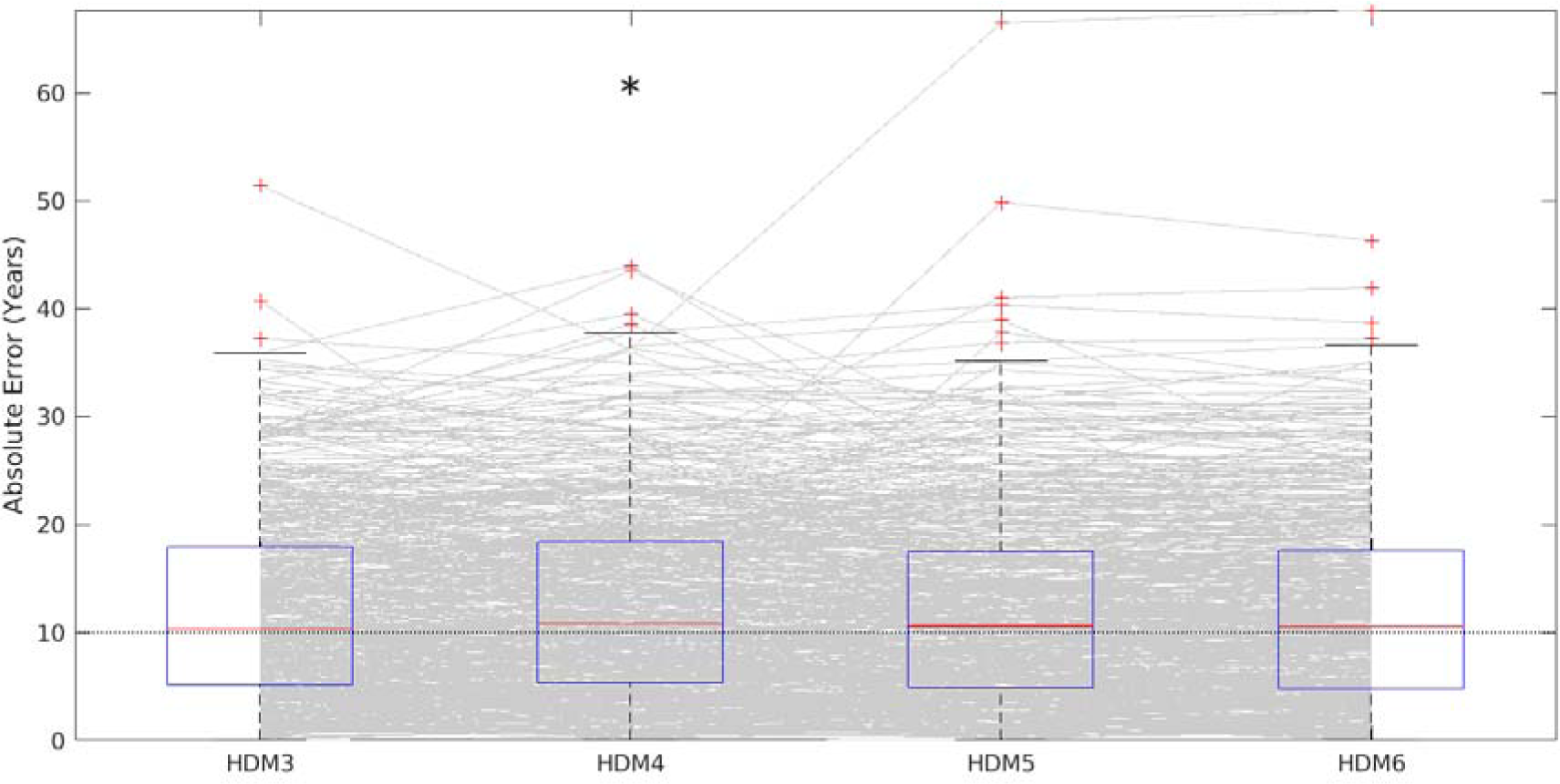
Cross-validated absolute error in predicting age, using the HDM model parameters combined across all ROIs, as the number of parameters increased. The HDM3 model is as in main paper; an HDM4 model with the additional vessel stiffness parameter *α*; an HDM5 model with the additional neurovascular feedback parameter *γ*; an HDM6 model with the additional oxygen extraction fraction *E_0_*. Each grey line corresponds to one participant. Superimposed on these lines are boxplots together with outliers (red crosses). An asterisk means that a sign-test revealed significantly different prediction than the simplest (HDM3) model. The average across ROIs of the median error was 10.3 years (HDM3), 10.8 years (HDM4), 10.6 years (HDM5) and 10.5 years (HDM6). Note this is using the parameter estimates before application of PEB.

**Supplementary Figure S8.**
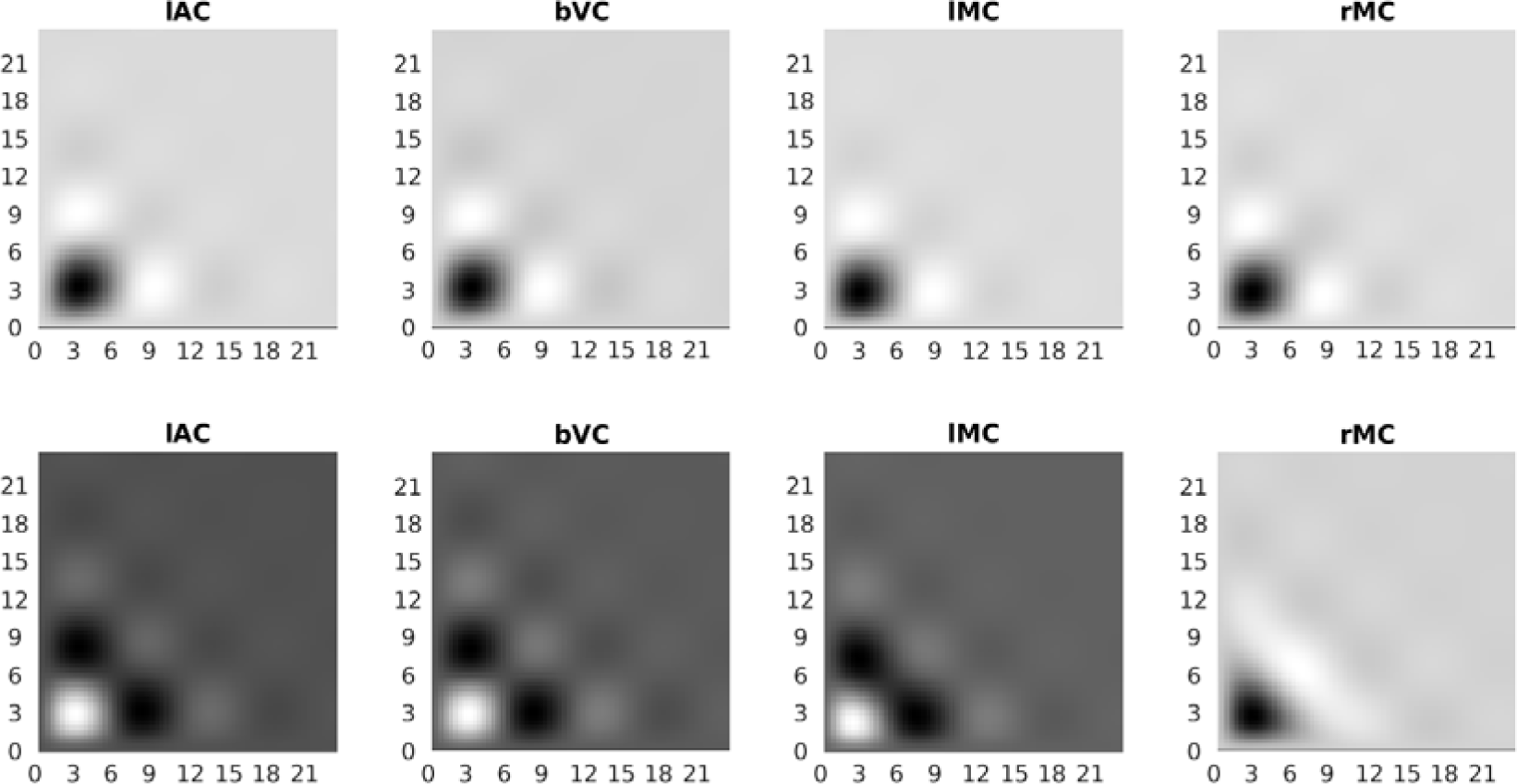
Second-order Volterra kernels for each ROI from the HDM3 model, capturing nonlinear effects of time between trial onsets (SOA; plotted on both axes). The top row shows the mean across participants, while the bottom row shows the effect of age. Note that in order to see effects (e.g, in rMC), the grayscale is optimised for each ROI and kernel separately (units are arbitrary). The dark region in the top row indicates under-additivity (saturation) for SOAs below ∼6s in all ROIs, while the white region shows super-additivity for SOAs from ∼6s to ∼10s, replicating previous findings from Friston et al. (1998). The maximal saturation varied for SOAs 2.6s-3.8s across ROIs, and was highly significant in all ROIs (T(636)>19.6, p<1e-16, though uncorrected for multiple comparisons). The bottom row shows that the under-additivity below ∼6s, as well as the super-additivity between 6-10s, is attenuated with age in all ROIs except rMC, where they are augmented with age. Taking the SOAs with the maximal effect in the top row (which does not bias the orthogonal age effect), the age-related effect was significant for all ROIs, two-tailed T(636)>2.52, p<.05. The percentage of variance from the two kernels that was explained by the second-order kernel had a median value across participants of 29.2%, 15.6%, 23.9% and 4.5% for lAC, bVC, lMC and rMC respectively. The proportion of that second-order variance related to age (linear and/or quadratic) was 12.1%, 6.1%, 4.5% and 1.3% respectively.

## Footnotes

1 except that orthogonalisation of regressors within each condition was switched off within spm_fMRI_design.m (see https://github.com/RikHenson/AgeingHRF)

2 The purpose of this study is to compare models of the effect of age on the HRF, so we want to define ROIs that show strong age effects in the first place. Though performing this ROI selection from fits of the FIR model biases the comparison of the FIR model with other models (as noted later), alternative definitions such as defining ROIs on the average effect across participants would not identify the right motor cortex, because it showed a particularly strong effect of age, while defining ROIs on anatomical criteria instead would be problematic because the age effects could reflect functional boundaries not respected by anatomy. In other words, our goal is not to identify (localise) brain regions that happen to show age effects in this paradigm, but rather explain the nature of those effects (in terms of HRF shape or underlying physiological parameters) wherever those age effects are strong.

3 The degrees of freedom in this model are so high (>20,000) that nearly every active voxel survives correction for multiple comparisons, hence the arbitrary threshold to define reasonably-sized ROIs.4 Trial-locked FIR effects that do not resemble a typical HRF are sometimes rejected as “non-hemodynamic”, but in this case, such an implicit model of the HRF would be better put explicitly into the GLM in the first place, e.g, by using a more constrained basis set. Nonetheless, if the neural activity associated with a trial is sustained (more than a second or so) and unknown in form, then a FIR basis set can be a useful “catch-all”.

4 Trial-locked FIR effects that do not resemble a typical HRF are sometimes rejected as “non-hemodynamic”, but in this case, such an implicit model of the HRF would be better put explicitly into the GLM in the first place, e.g, by using a more constrained basis set. Nonetheless, if the neural activity associated with a trial is sustained (more than a second or so) and unknown in form, then a FIR basis set can be a useful “catch-all”.

5 Note that, unlike the other 3 models, the NLF4 approach is fit to the FIR parameters (i.e, peristimulus time), rather than the original fMRI timeseries. One could take the best-fitting HRF shape from such nonlinear fitting of the FIR parameters, and insert it back into the first-level GLM for the fMRI timeseries, in order to estimate a single scaling (amplitude) parameter. This would enable inference to be performed on a single parameter, while allowing for variability in the HRF shape across ROIs and/or across participants. This has the potential to be more robust than re-inserting the FIR estimate of HRF shape for a given participant and ROI, if the latter is noisy, owing to the effective “regularisation” of the HRF by the average shape across participants and/or ROIs (similar to the formal hierarchical model used for HDM below). Nonetheless, care would be needed for regions like rMC that do not show a consistent HRF shape across participants (see Results).

6 The autoregulatory feedback parameter may not have a direct physiological correlate at the vascular level. Indeed, it has been proposed that neurovascular coupling can be modelled as a purely feedforward system, in conjunction with inhibitory neural activity (Havlicek et al., 2015). In the absence of a neural model, the purpose of the feedback parameter here is simply to enable damped oscillatory dynamics on a physiologically plausible timescale. See Appendix 5 of Zeidman et al. (2019) for details.

7 Statistical parametric maps for the stimulus-locked and response-locked group FIR models were similar, as would be expected since the median across participants of their median reaction time (RT) across trials was 282ms (range = 182ms - 679ms), which is short relative to the time-constants of the BOLD response. Nonetheless, this variability in RTs (across both trials and participants) is important for latency analyses below.

8 The degrees of freedom in this model are so high (>20,000) that nearly every active voxel would survive correction for multiple comparisons, so the statistical and extent thresholds were chosen simply to obtain reasonably-sized ROIs.

9 Note that the slight “sawtooth” modulation across alternating FIR bins (particularly apparent in tail of HRFs) reflects the differential sampling across the run owing to the non-integer ratio of bin-width relative to TR, such that even-numbered bins tended to be sampled from trials later in the run than odd-numbered bins.

10 We did try a fifth model in which the gamma function for the peak and the gamma function for the undershoot were separate basis functions (along with their temporal derivatives, i.e. 4 basis functions in total), but this model did not perform as well (e.g., in the cross-validated prediction of age reported later). Also note that, according to the HDM model, the peak and undershoot are highly coupled, since they are part of the same damped oscillation in response to a brief burst of neural activity.

11 We did try adding second-order expansions of HDM parameters (including interaction terms), as well as more advanced ML techniques (such as gradient boosting), but these did not show better cross-validated prediction than standard (unregularised) multiple regression (indeed, SHAP analyses showed linear terms are sufficient).

12 One could also enter such independent measures as additional covariates in the PEB model (as could higher-order polynomial expansions of age, in case there are nonlinear effects of age on the HDM parameters). However, given the high correlation of these vascular/neural factors with age, here we sought instead to test whether such measures mediated the age effects that we already found.

13 The more constrained nature of the Can3 basis set also prevents false detection of trial-locked effects that are implausible as hemodynamic responses, such as “activity” in the eyeballs for the first FIR time bin only, which reflects stimulus-driven eye-motion instead.

## References

Abdelkarim, D., Zhao, Y., Turner, M. P., Sivakolundu, D. K., Lu, H., & Rypma, B. (2019). A neural-vascular complex of age-related changes in the human brain: Anatomy, physiology, and implications for neurocognitive aging. Neuroscience and Biobehavioral Reviews, 107(December 2018), 927–944. 10.1016/j.neubiorev.2019.09.005

Aizenstein, H. J., Clark, K. A., Butters, M. A., Cochran, J., Stenger, V. A., Meltzer, C. C., Reynolds, C. F., & Carter, C. S. (2004). The BOLD hemodynamic response in healthy aging. Journal of Cognitive Neuroscience, 16(5), 786–793. 10.1162/089892904970681

Birn, R. M., Saad, Z. S., & Bandettini, P. A. (2001). Spatial Heterogeneity of the Nonlinear Dynamics in the FMRI BOLD Response. NeuroImage, 14(4), 817–826. 10.1006/nimg.2001.0873

Buračas, G. T., & Boynton, G. M. (2002). Efficient design of event-related fMRI experiments using m-sequences. NeuroImage, 16(3 I), 801–813. 10.1006/nimg.2002.1116

Buxton, R. B., Wong, E. C., & Frank, L. R. (1998). Dynamics of blood flow and oxygenation changes during brain activation: The balloon model. Magnetic Resonance in Medicine, 39(6), 855– 864. 10.1002/mrm.1910390602

Calhoun, V. D., Stevens, M. C., Pearlson, G. D., & Kiehl, K. A. (2004). fMRI analysis with the general linear model: Removal of latency-induced amplitude bias by incorporation of hemodynamic derivative terms. NeuroImage, 22(1), 252–257. 10.1016/j.neuroimage.2003.12.029

Cignetti, F., Salvia, E., Anton, J. L., Grosbras, M. H., & Assaiante, C. (2016). Pros and Cons of Using the Informed Basis Set to Account for Hemodynamic Response Variability with Developmental Data. Frontiers in Neuroscience, 10(JUL), 322. 10.3389/FNINS.2016.00322

Cusack, R., Vicente-Grabovetsky, A., Mitchell, D. J., Wild, C. J., Auer, T., Linke, A. C., & Peelle, J. E. (2015). Automatic analysis (aa): Efficient neuroimaging workflows and parallel processing using Matlab and XML. Frontiers in Neuroinformatics, 8(JAN), 1–13. 10.3389/fninf.2014.00090

Dale, A. M., & Buckner, R. L. (1997). Selective Averaging of Rapidly Presented Individual Trials Using fMRI. Hum. Brain Mapping, 5, 329–340. 10.1002/(SICI)1097-0193(1997)5:5

D’Esposito, M., Zarahn, E., Aguirre, G. K., & Rypma, B. (1999). The effect of normal aging on the coupling of neural activity to the bold hemodynamic response. NeuroImage, 10(1), 6–14. 10.1006/nimg.1999.0444

Friston, K. J., Fletcher, P., Josephs, O., Holmes, A., Rugg, M. D., & Turner, R. (1998). Event-Related fMRI: Characterizing Differential Responses. 40(7), 30–40.

Friston, K. J., Harrison, L., & Penny, W. (2003). Dynamic causal modelling. NeuroImage, 19(4), 1273– 1302. 10.1016/S1053-8119(03)00202-7

Friston, K. J., Josephs, O., Rees, G., & Turner, R. (1998). Nonlinear event-related responses in fMRI. Magnetic Resonance in Medicine, 39(1), 41–52. 10.1002/mrm.1910390109

Friston, K. J., Mechelli, A., Turner, R., & Price, C. J. (2000). Nonlinear Responses in fMRI: The Balloon Model, Volterra Kernels, and Other Hemodynamics. NeuroImage, 12(4), 466–477. 10.1006/NIMG.2000.0630

Fuhrmann, D., Nesbitt, D., Shafto, M., Rowe, J. B., Price, D., Gadie, A., Tyler, L. K., Brayne, C., Bullmore, E. T., Calder, A. C., Cusack, R., Dalgleish, T., Duncan, J., Henson, R. N., Matthews, F. E., Marslen-Wilson, W. D., Rowe, J. B., Shafto, M. A., Campbell, K., … Kievit, R. A. (2019). Strong and specific associations between cardiovascular risk factors and white matter micro- and macrostructure in healthy aging. Neurobiology of Aging, 74, 46–55. 10.1016/j.neurobiolaging.2018.10.005

Graff, B. J., Harrison, S. L., Payne, S. J., & El-Bouri, W. K. (2023). Regional Cerebral Blood Flow Changes in Healthy Ageing and Alzheimer’s Disease: A Narrative Review. Cerebrovascular Diseases, 52(1), 11–20. 10.1159/000524797

Grinband, J., Steffener, J., Razlighi, Q. R., & Stern, Y. (2017). BOLD neurovascular coupling does not change significantly with normal aging. Human Brain Mapping, 38(7), 3538–3551. 10.1002/hbm.23608

Grubb, R. L., Raichle, M. E., Eichling, J. O., & Ter-Pogossian, M. M. (1974). The Effects of Changes in PaCO2 Cerebral Blood Volume, Blood Flow, and Vascular Mean Transit Time. Stroke, 5(5), 630–639. 10.1161/01.STR.5.5.630

Handwerker, D. A., Gazzaley, A., Inglis, B. A., & D’Esposito, M. (2007). Reducing vascular variability of fMRI data across aging populations using a breathholding task. Human Brain Mapping, 28(9), 846–859. 10.1002/hbm.20307

Handwerker, D. A., Gonzalez-Castillo, J., D’Esposito, M., & Bandettini, P. A. (2012). The continuing challenge of understanding and modeling hemodynamic variation in fMRI. NeuroImage, 62(2), 1017–1023. 10.1016/j.neuroimage.2012.02.015

Havlicek, M., Roebroeck, A., Friston, K., Gardumi, A., Ivanov, D., & Uludag, K. (2015). Physiologically informed dynamic causal modeling of fMRI data. NeuroImage, 122, 355–372. 10.1016/j.neuroimage.2015.07.078

Havlicek, M., Roebroeck, A., Friston, K. J., Gardumi, A., Ivanov, D., & Uludag, K. (2017). On the importance of modeling fMRI transients when estimating effective connectivity: A dynamic causal modeling study using ASL data. NeuroImage, 155, 217–233. 10.1016/j.neuroimage.2017.03.017

Henson, R. (2004). Analysis of fMRI Timeseries: Linear Time-Invariant Models, Event-related fMRI and Optimal Experimental Design. In Human Brain Function (Second Edition) (pp. 793–822). 10.1006/nimg.1995.1023

Henson, R. N. A., Price, C. J., Rugg, M. D., Turner, R., & Friston, K. J. (2002). Detecting latency differences in event-related BOLD responses: Application to words versus nonwords and initial versus repeated face presentations. NeuroImage, 15(1), 83–97. 10.1006/nimg.2001.0940

Josephs, O., & Henson, R. N. A. (1999). Event-related functional magnetic resonance imaging: Modelling, inference and optimization. Philosophical Transactions of the Royal Society of London. Series B: Biological Sciences, 354(1387), 1215–1228. 10.1098/rstb.1999.0475

King, D. L. O., Henson, R. N., Kievit, R., Wolpe, N., Brayne, C., Tyler, L. K., Rowe, J. B., Cam-CAN, Bullmore, E. T., Calder, A. C., Cusack, R., Dalgleish, T., Duncan, J., Matthews, F. E., Marslen-Wilson, W. D., Shafto, M. A., Campbell, K., Cheung, T., Davis, S., … Tsvetanov, K. A. (2023). Distinct components of cardiovascular health are linked with age-related differences in cognitive abilities. Scientific Reports, 13(1), 978. 10.1038/s41598-022-27252-1

Knights, E., Morcom, A. M., & Henson, R. N. (2021). Does hemispheric asymmetry reduction in older adults in motor cortex reflect compensation? Journal of Neuroscience, 41(45), 9361–9378. 10.1523/JNEUROSCI.1111-21.2021

Kruggel, F., & Yves Von Cramon, D. (1999). Modeling the Hemodynamic Response in Single-Trial Functional MRI Experiments. Magnetic Resonance in Medicine, 42, 787–797. 10.1002/(SICI)1522-2594(199910)42:4

Lindquist, M. A., Meng Loh, J., Atlas, L. Y., & Wager, T. D. (2009). Modeling the Hemodynamic Response Function in fMRI: Efficiency, Bias and Mis-modeling. Neuroimage, 45(1 Suppl), S187. 10.1016/J.NEUROIMAGE.2008.10.065

Marreiros, A. C., Kiebel, S. J., & Friston, K. J. (2008). Dynamic causal modelling for fMRI: A two-state model. NeuroImage, 39(1), 269–278. 10.1016/j.neuroimage.2007.08.019

Mattay, V. S., & Weinberger, D. R. (1999). Organization of the human motor system as studied by functional magnetic resonance imaging. European Journal of Radiology, 30(2), 105–114. 10.1016/S0720-048X(99)00049-2

Mayhew, S. D., Coleman, S. C., Mullinger, K. J., & Can, C. (2022). Across the adult lifespan the ipsilateral sensorimotor cortex negative BOLD response exhibits decreases in magnitude and spatial extent suggesting declining inhibitory control. NeuroImage, 253. 10.1016/j.neuroimage.2022.119081

Price, D., Tyler, L. K., Neto Henriques, R., Campbell, K. L., Williams, N., Treder, M. S., Taylor, J. R., Henson, R. N. A., Brayne, C., Bullmore, E. T., Calder, A. C., Cusack, R., Dalgleish, T., Duncan, J., Matthews, F. E., Marslen-Wilson, W. D., Rowe, J. B., Shafto, M. A., Cheung, T., … Villis, L. (2017). Age-related delay in visual and auditory evoked responses is mediated by white-and grey-matter differences. Nature Communications, 8(May 2016). 10.1038/ncomms15671

Richter, W., & Richter, M. (2003). The shape of the fMRI BOLD response in children and adults changes systematically with age. NeuroImage, 20(2), 1122–1131. 10.1016/S1053-8119(03)00347-1

Shafto, M. A., Tyler, L. K., Dixon, M., Taylor, J. R., Rowe, J. B., Cusack, R., Calder, A. J., Marslen-Wilson, W. D., Duncan, J., Dalgleish, T., Henson, R. N., Brayne, C., Bullmore, E., Campbell, K., Cheung, T., Davis, S., Geerligs, L., Kievit, R., McCarrey, A., … Matthews, F. E. (2014). The Cambridge Centre for Ageing and Neuroscience (Cam-CAN) study protocol: A cross-sectional, lifespan, multidisciplinary examination of healthy cognitive ageing. BMC Neurology, 14(1). 10.1186/S12883-014-0204-1

Shrout, P. E., & Bolger, N. (2002). Mediation in experimental and nonexperimental studies: New procedures and recommendations. Psychological Methods, 7(4), 422–445.

Stephan, K. E., Weiskopf, N., Drysdale, P. M., Robinson, P. A., & Friston, K. J. (2007). Comparing hemodynamic models with DCM. NeuroImage, 38(3), 387–401. 10.1016/J.NEUROIMAGE.2007.07.040

Stiernman, L., Grill, F., McNulty, C., Bahrd, P., Panes Lundmark, V., Axelsson, J., Salami, A., & Rieckmann, A. (2023). Widespread fMRI BOLD signal overactivations during cognitive control in older adults are not matched by corresponding increases in fPET glucose metabolism. The Journal of Neuroscience, 43(14), JN--RM--1331--22. 10.1523/jneurosci.1331-22.2023

Tak, Y. W., Knights, E., Henson, R., & Zeidman, P. (2021). Ageing and the ipsilateral m1 bold response: A connectivity study. Brain Sciences, 11(9). 10.3390/BRAINSCI11091130/S1

Taoka, T., Iwasaki, S., Uchida, H., Fukusumi, A., Nakagawa, H., Kichikawa, K., Takayama, K., Yoshioka, T., Takewa, M., & Ohishi, H. (1998). Age Correlation of the Time Lag in Signal Change on EPI-fMRI. Journal of Computer Assisted Tomography, 22(4). https://journals.lww.com/jcat/Fulltext/1998/07000/Age_Correlation_of_the_Time_Lag_in_Signal_Change.2.aspx

Taylor, J. R., Williams, N., Cusack, R., Auer, T., Shafto, M. A., Dixon, M., Tyler, L. K., Cam-CAN, & Henson, R. N. (2017). The Cambridge Centre for Ageing and Neuroscience (Cam-CAN) data repository: Structural and functional MRI, MEG, and cognitive data from a cross-sectional adult lifespan sample. NeuroImage, 144(Pt B), 262–269. 10.1016/J.NEUROIMAGE.2015.09.018

Tsvetanov, K. a., Henson, R. N. a., Tyler, L. K., Razi, A., Geerligs, L., Ham, T. E., & Rowe, J. B. (2016). Extrinsic and Intrinsic Brain Network Connectivity Maintains Cognition across the Lifespan Despite Accelerated Decay of Regional Brain Activation. Journal of Neuroscience, 36(11), 3115–3126. 10.1523/JNEUROSCI.2733-15.2016

Tsvetanov, K. A., Henson, R. N. A., & Rowe, J. B. (2021). Separating vascular and neuronal effects of age on fMRI BOLD signals: Neurovascular ageing. Philosophical Transactions of the Royal Society B: Biological Sciences, 376(1815). 10.1098/rstb.2019.0631

Tsvetanov, K. A., Henson, R. N. A., Tyler, L. K., Davis, S. W., Shafto, M. A., Taylor, J. R., Williams, N., Cam-CAN, & Rowe, J. B. (2015). The effect of ageing on fMRI: Correction for the confounding effects of vascular reactivity evaluated by joint fMRI and MEG in 335 adults. Human Brain Mapping, 36(6), 2248. 10.1002/hbm.22768

Vazquez, A. L., Fukuda, M., & Kim, S. G. (2018). Inhibitory Neuron Activity Contributions to Hemodynamic Responses and Metabolic Load Examined Using an Inhibitory Optogenetic Mouse Model. Cerebral Cortex, 28(11), 4105–4119. 10.1093/CERCOR/BHY225

Ward, N. S., Swayne, O. B. C., & Newton, J. M. (2008). Age-dependent changes in the neural correlates of force modulation: An fMRI study. Neurobiology of Aging, 29(9), 1434–1446. 10.1016/j.neurobiolaging.2007.04.017

West, K. L., Zuppichini, M. D., Turner, M. P., Sivakolundu, D. K., Zhao, Y., Abdelkarim, D., Spence, J. S., & Rypma, B. (2019). BOLD hemodynamic response function changes significantly with healthy aging. NeuroImage, 188, 198–207. 10.1016/J.NEUROIMAGE.2018.12.012

Wright, M. E., & Wise, R. G. (2018). Can Blood Oxygenation Level Dependent Functional Magnetic Resonance Imaging Be Used Accurately to Compare Older and Younger Populations? A Mini Literature Review. Frontiers in Aging Neuroscience, 10. 10.3389/FNAGI.2018.00371

Zeidman, P., Jafarian, A., Seghier, M. L., Litvak, V., Cagnan, H., Price, C. J., & Friston, K. J. (2019). A guide to group effective connectivity analysis, part 2: Second level analysis with PEB. NeuroImage, 200, 12–25. 10.1016/J.NEUROIMAGE.2019.06.032

## References

Buxton RB, Wong EC and Frank LR (1998) Dynamics of blood flow and oxygenation changes during brain activation: The balloon model. Magnetic Resonance in Medicine 39(6): 855–864. DOI: 10.1002/mrm.1910390602.

Dubeau S, Ferland G, Gaudreau P, et al. (2011) Cerebrovascular hemodynamic correlates of aging in the Lou / c rat: A model of healthy aging. NeuroImage 56(4). Elsevier Inc.: 1892–1901. DOI: 10.1016/j.neuroimage.2011.03.076.

Friston KJ, Mechelli A, Turner R, et al. (2000) Nonlinear Responses in fMRI: The Balloon Model, Volterra Kernels, and Other Hemodynamics. NeuroImage 12(4). Academic Press: 466–477. DOI: 10.1006/NIMG.2000.0630.

Friston KJ, Litvak V, Oswal A, et al. (2016) Bayesian model reduction and empirical Bayes for group (DCM) studies. NeuroImage 128. The Authors: 413–431. DOI: 10.1016/j.neuroimage.2015.11.015.

Grubb RL, Raichle ME, Eichling JO, et al. (1974) The Effects of Changes in PaCO2 Cerebral Blood Volume, Blood Flow, and Vascular Mean Transit Time. Stroke 5(5). Lippincott Williams & Wilkins: 630–639. DOI: 10.1161/01.STR.5.5.630.

Havlicek M, Roebroeck A, Friston K, et al. (2015) Physiologically informed dynamic causal modeling of fMRI data. NeuroImage 122. Elsevier B.V.: 355–372. DOI: 10.1016/j.neuroimage.2015.07.078.

Heinzle J, Koopmans PJ, den Ouden HEM, et al. (2016) A hemodynamic model for layered BOLD signals. NeuroImage 125. Elsevier Inc.: 556–570. DOI: 10.1016/j.neuroimage.2015.10.025.

Hua J, Liu P, Kim T, et al. (2019) MRI techniques to measure arterial and venous cerebral blood volume. NeuroImage 187. Neuroimage: 17–31. DOI: 10.1016/J.NEUROIMAGE.2018.02.027.

Leenders KL, Perani D, Lammertsma AA, et al. (1990) Cerebral blood flow, blood volume and oxygen utilization. Normal values and effect of age. Brain: a journal of neurology 113 (Pt 1)(1). Brain: 27–47. DOI: 10.1093/BRAIN/113.1.27.

Leung TS, Tachtsidis I, Tisdall MM, et al. (2008) Estimating a modified Grubb’s exponent in healthy human brains with near infrared spectroscopy and transcranial Doppler. Physiological Measurement 30(1). IOP Publishing: 1. DOI: 10.1088/0967-3334/30/1/001.

Peng SL, Dumas JA, Park DC, et al. (2014) Age-related increase of resting metabolic rate in the human brain. NeuroImage 98. Neuroimage: 176–183. DOI: 10.1016/J.NEUROIMAGE.2014.04.078.

Stephan KE, Weiskopf N, Drysdale PM, et al. (2007) Comparing hemodynamic models with DCM. NeuroImage 38(3). Neuroimage: 387–401. DOI: 10.1016/J.NEUROIMAGE.2007.07.040.

Uludag K, Müller-bierl B and Kâmil U (2009) An integrative model for neuronal activity-induced signal changes for gradient and spin echo functional imaging. NeuroImage 48: 150–165. DOI: 10.1016/j.neuroimage.2009.05.051.

Zeidman P, Jafarian A, Seghier ML, et al. (2019) A guide to group effective connectivity analysis, part 2: Second level analysis with PEB. NeuroImage 200. Academic Press: 12–25. DOI: 10.1016/J.NEUROIMAGE.2019.06.032.

