## Supplementary material for "Evaluating models of the ageing BOLD response": All Sup Mat

- The neurovascular parameter $\gamma$ is the rate constant controlling the feedback from blood flow. Together with the decay parameter $\kappa$, the two produce a damped oscillation of the vasodilatory signal. However the primary effect of decreasing $\gamma$ is to delay the HRF, which is similar to the primary effect of decreasing the haemodynamic transit rate $1/\tau_h$ (Supplementary Figure S1), resulting in a high negative correlation between them (Supplementary Figure S2). For the HDM3 model, we therefore fixed $\gamma$ here based on its prior expected value of 0.41Hz from Friston et al. (2000) (though allowed it to vary in the HDM5 model below). This does mean, however, that changes in neurovascular feedback could also reflect changes in haemodynamic transit rate.

- We adjusted the values of parameters $\epsilon_h$, $r_0$ and $\vartheta_0$ to correspond to the MRI field strength for the CamCAN data, where $B_0 = 3T$. In previous implementations of this model, the ratio of intra-vascular to extra-vascular signal, $\epsilon_h$, was estimated from the data, reflecting uncertainty about its value in the literature (Stephan et al., 2007). Here, we decided to fix its value for stability, as large values of $\epsilon_h$ can induce phase transitions that give rise to unrealistic BOLD responses. Following Havlicek et al., 2015, we used the expected value of $\epsilon_h = 0.44$, based on the ranges of T2* relaxation rates (for further detail, see the MATLAB script *Heinzle_epsilon_derivation.m* in the HDM Toolbox, https://github.com/pzeidman/HDM-toolbox). We set the value of $r_0 = 110s^{-1}$ and $\vartheta_0 = 28.265B_0 = 31.27s^{-1}$, following Heinzle et al. (2016), who drew on the results of Uludag et al. (2009).

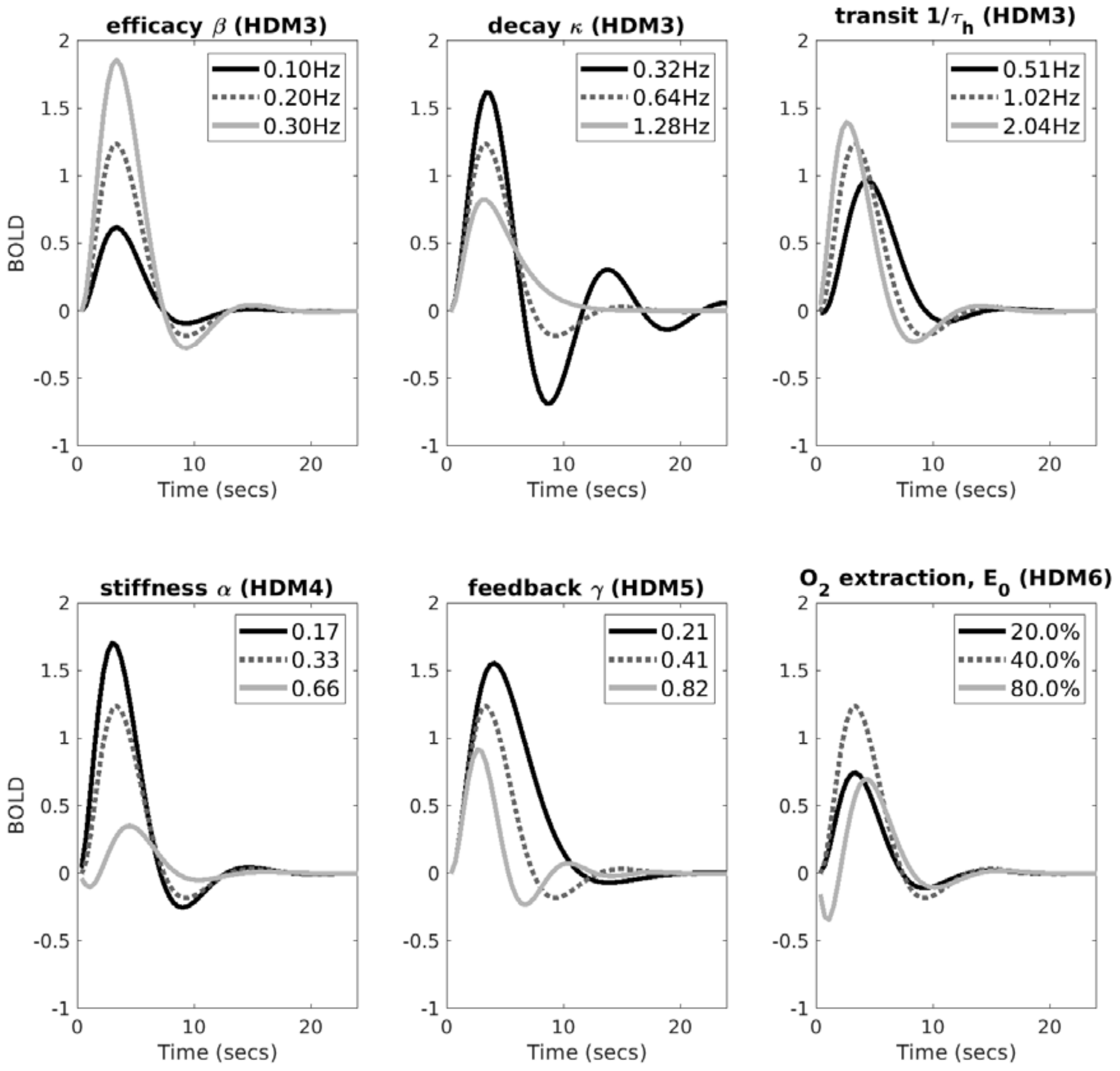


Supplementary Figure S1. Each plot shows the predicted BOLD response under different values of a parameter, indicated by the title. The central parameter value (producing the dotted line) reflects the prior expectation; the dark solid line and light solid line reflect smaller or larger values respectively (the values of all parameters other than the one varied in a plot were fixed at their prior expectation, except neural efficacy, β, which was set to 0.2).

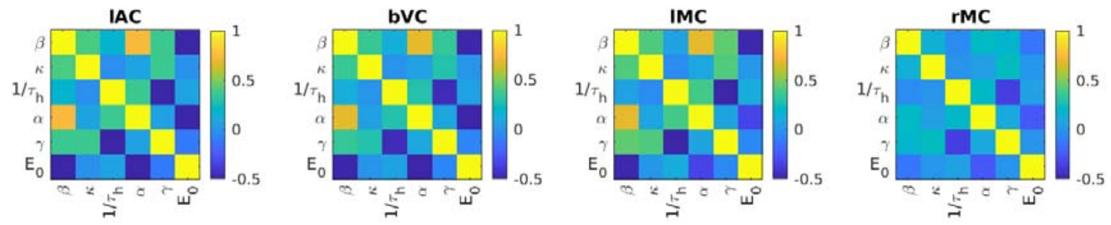


Supplementary Figure S2. The mean across participants of the posterior correlation between HDM parameters after fitting a 6-parameter HDM model (HDM6) to the data in the paper. Apart from rMC, note that the other ROIs have a large positive correlation between neural efficacy $\beta$ and vascular stiffness $\alpha$, and large negative correlations between neurovascular feedback $\gamma$ and haemodynamic transit rate $1/\tau_h$, and between oxygen extraction $E_0$ and $\beta$.

A summary of the prior expected value and variance for the three free parameters in the main HDM3 model used in the paper are shown in Supplementary Table 1. Note that, in order to enforce positivity constraints on two of these parameters, new parameters $l_\kappa$ and $l_{\tau_h}$ are introduced, which are log versions of decay rate $\kappa$ and transit rate, $1/\tau_h$, respectively. The scaling values of 0.64 and 1.02 are taken from (Friston et al., 2000). The neural efficacy parameter $\beta$ is not constrained to be positive so is untransformed.

Additionally, in order to enforce positivity constraints on the states $f_{in}$, $v$ and $q$, each variable is log-transformed, requiring the equations to be supplemented as follows:

$$\frac{d \ln f_{in}}{dt} = \frac{\dot{f}_{in}}{f_{in}} \quad (9)$$

$$\frac{d \ln v}{dt} = \frac{\dot{v}}{v} \quad (10)$$

$$\frac{d \ln q}{dt} = \frac{\dot{q}}{q} \quad (11)$$

**Supplementary Table 1: Free parameters of HDM3 model**

| Model | Parameter | Compartment | Prior expectation | Prior variance | Parameterisation |
|---|---|---|---|---|---|
| HDM3 | $\beta$ | Neural | 0 | 1 | - |
| HDM3 | $l_\kappa$ | Neurovascular (CBF) | 0 | 1/32 | $\kappa = 0.64s^{-1} \cdot \exp(l_k)$ |
| HDM3 | $l_{\tau_h}$ | Vascular (CBV) | 0 | 1/32 | $\frac{1}{\tau_h} = 1.02s^{-1} \cdot \exp(l_{\tau_h})$ |

The HDM models with 4, 5 and 6 free parameters (used later in Sup Mat) included the values in Supplementary Table 1, plus the one or more of rows of Supplementary Table 2.

**Supplementary Table 2: Additional free parameters of HDM4, HDM5 and HDM6 models**

| Model | Parameter | Compartment | Prior expectation | Prior variance | Parameterisation |
|---|---|---|---|---|---|
| HDM4 | $l_\alpha$ | Vascular | 0 | 1/32 | $\alpha = 0.33 \cdot \exp(l_\alpha)$ |
| HDM5 | $l_\gamma$ | Neurovascular | 0 | 1/32 | $\gamma = 0.41s^{-1} \cdot \exp(l_\gamma)$ |
| HDM6 | $l_{E_0}$ | Vascular | 0 | 1/32 | $E_0 = 0.40 \cdot \exp(l_{E_0})$ |

### Group-level model using Parametric Empirical Bayes (PEB)

For the PEB estimation of a group-level model (Zeidman et al., 2019), parameter estimates from all $n$ participants are concatenated into a vector of random variables, $\theta^{(1)} = (\theta_1, \theta_2 \dots \theta_n)$. Between-participant effects were then modelled using a general linear model:

$$\theta^{(1)} = X\theta^{(2)} + \epsilon \tag{12}$$

The design matrix $X$ included regressors for the effect of each of two covariates (mean over participants and age) on each of $P = 3$ haemodynamic parameters ($\beta, l_\kappa, l_{\tau_h}$). This design matrix can be written formally as:

$$X = X_B \otimes I_P \tag{13}$$

where between-participants design matrix $X_B$ contained two columns: a column of ones (to model the average haemodynamic parameters over participants) and the z-scored age of each participant. This was replicated over the three parameters by taking the Kronecker tensor product ⊗ with the identity matrix of dimension $P$.

Unexplained between-participants variability $\epsilon$ was modelled using a covariance component model, with a separate I.I.D. precision component for each of the three haemodynamic parameters, allowing each type of parameter to have a separate level of between-participants variability.

The parameters $\theta^{(2)}$ and free energy $F$ of the PEB model were estimated using standard routines in the SPM software. We then tested the evidence for the presence versus absence of each covariate on each haemodynamic parameter using an automated procedure. This was an automatic search that iteratively pruned mixtures of group-level parameters $\theta^{(2)}$ from the PEB model, where doing so did not reduce the free energy. This was performed using an analytic approach called Bayesian Model Reduction (Friston et al., 2016).

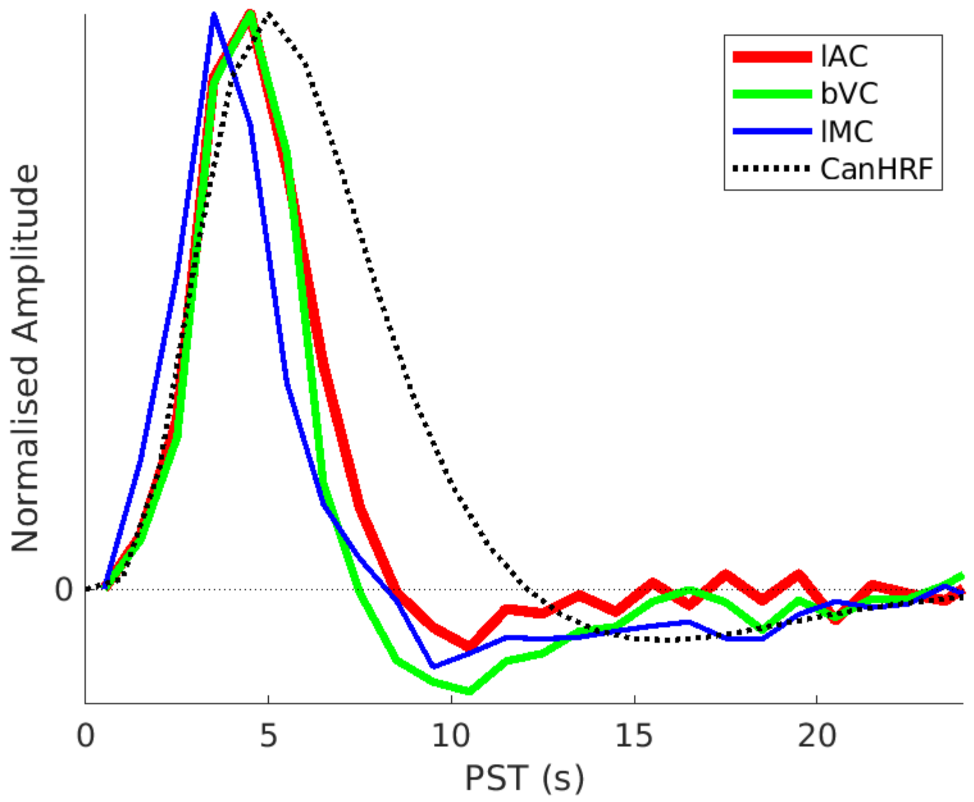


Supplementary Figure S3. Mean across all participants of the FIR fits for lAC, bVC and lMC (solid lines), along with SPM's canonical HRF (dotted line). The rMC is not shown because it varied so much with age (see main text). Peak amplitude is matched by scaling by the maximum positive value. The lAC and bVC data are from the stimulus-locked model, while lMC is from the response-locked model.

SPM's canonical HRF is more dispersed than the FIR fits (delayed peak and undershoot). This may not have been noticed before because early studies generating this canonical shape (Friston et al., 2000) did not allow for slice-timing delays. So a delay of TR/2 (typically ~1s) could partly explain this mismatch. Here we did correct the data for different slice-times and were careful to synchronise the GLM (for FIR32 and Can3 basis sets) with the reference slice used for slice-timing. Grinwald et al. (2017) still observed deviations despite slice-timing, though ones that derivatives should be able to accommodate.

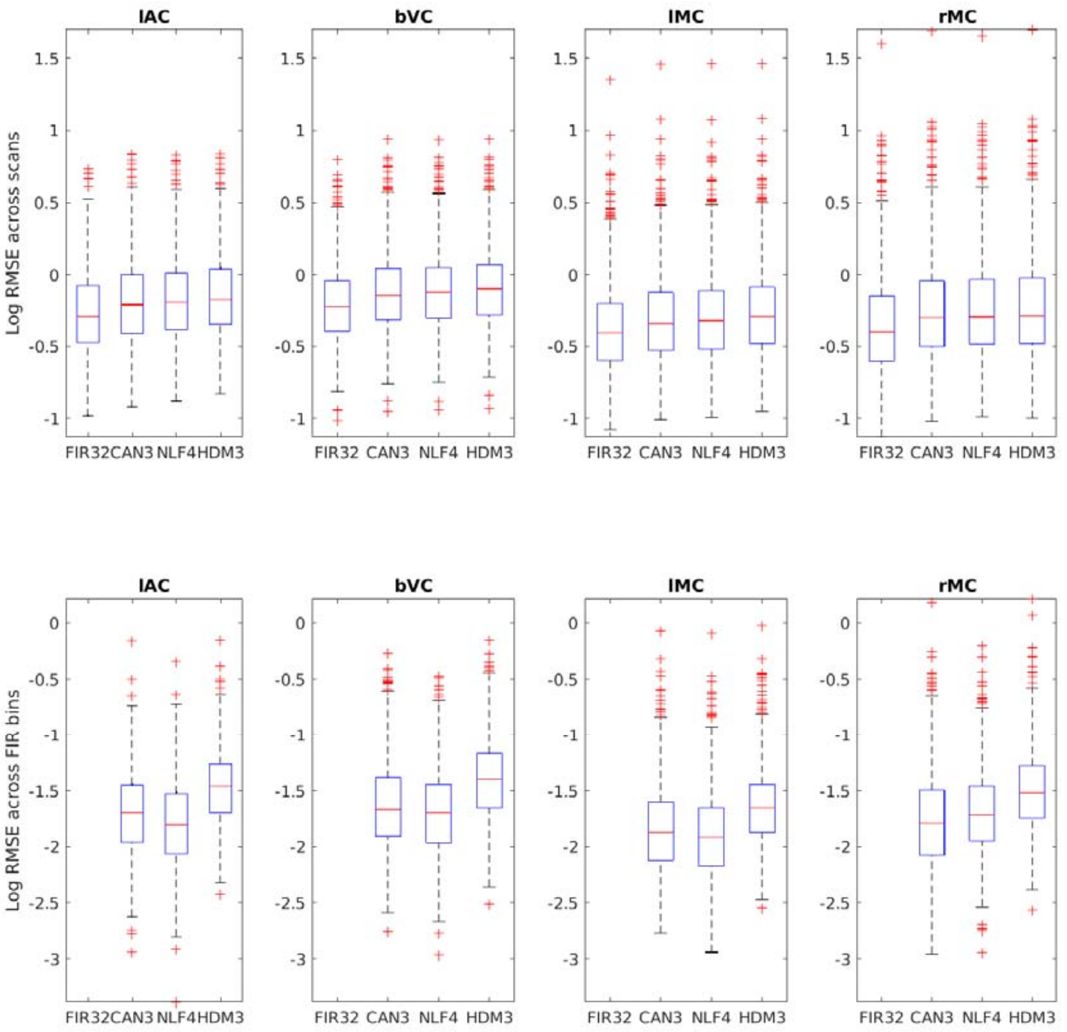


Supplementary Figure S4a. Boxplots (of log) of Root of Mean of Squared Error (RMSE) across scans (top row) or across FIR bins (bottom row) for each ROI and model. Note there is statistical circularity for the FIR model, since the same model was used to define the ROIs in the first place, but its inclusion here at least provides a lower bound, albeit biased, with which to compare the other models. Note also that there can be no RMSE for the FIR32 model in bottom row, and in the top row, the RMSE across scans for the NLF model was calculated by re-inserting the participant- and ROI-specific fitted HRF into each participant's first-level GLM (hence only 1 effective degree of freedom). The lAC and bVC data are from the stimulus-locked model, while lMC and rMC are from the response-locked model. Note that these measures of model fit ignore differences in model complexity, and so are not as good for generalisation to new data as the cross-validated error in Figure 9.

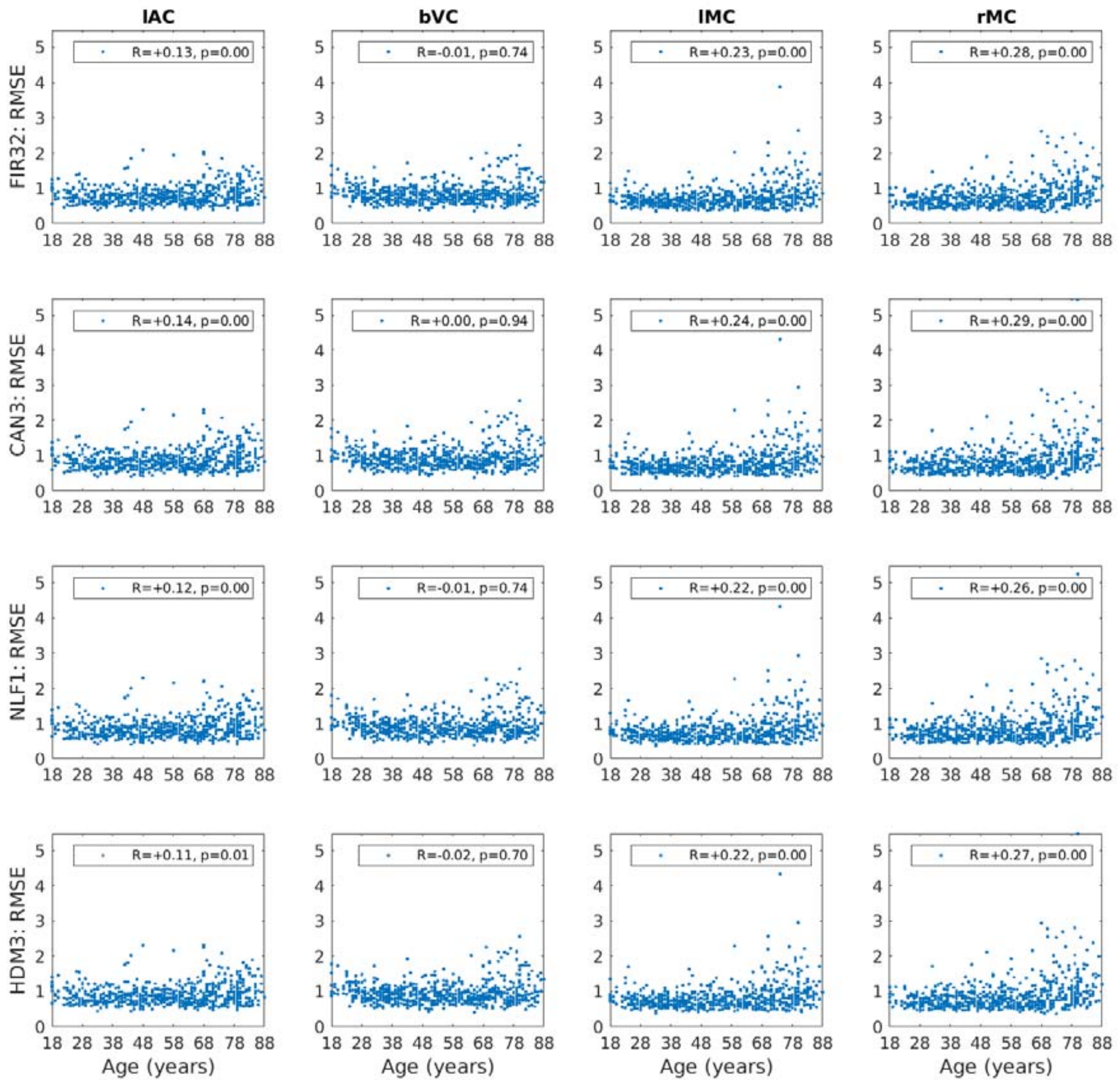


Supplementary Figure S4b. Square-root of Mean of Squared residuals (RMSE) across scans for each model and for each ROI, as a function of participant's age, together with Spearman correlation R- and p-values. The lAC and bVC data are from the stimulus-locked model, while lMC and rMC are from the response-locked model. Note that the RMSE for the NLF model was calculated by re-inserting the participant- and ROI-specific fitted HRF into each participant's first-level GLM (hence only 1 effective degree of freedom).

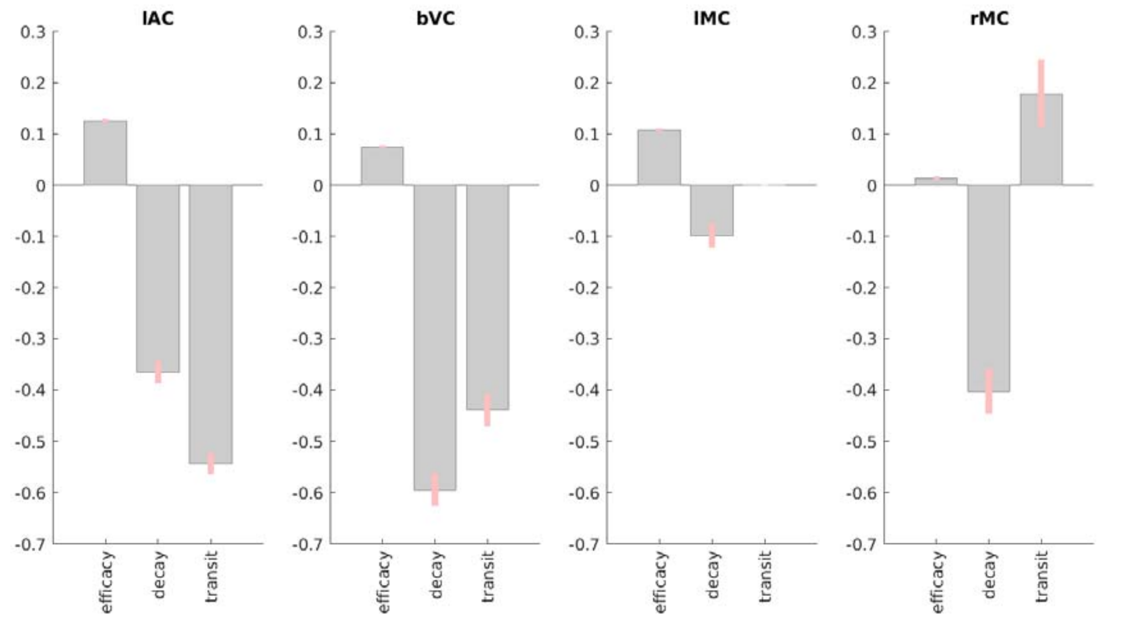


Supplementary Figure S5. Results of PEB BMR on HDM parameters for each ROI for the mean across participants, in terms of deviation of each parameter from its prior expectation. Grey bars show posterior expectation with 90% credible interval in pink; missing bars are parameters that BMR has removed as unnecessary (in terms of maximising evidence for PEB model). For decay and transit parameters, units are log deviations.

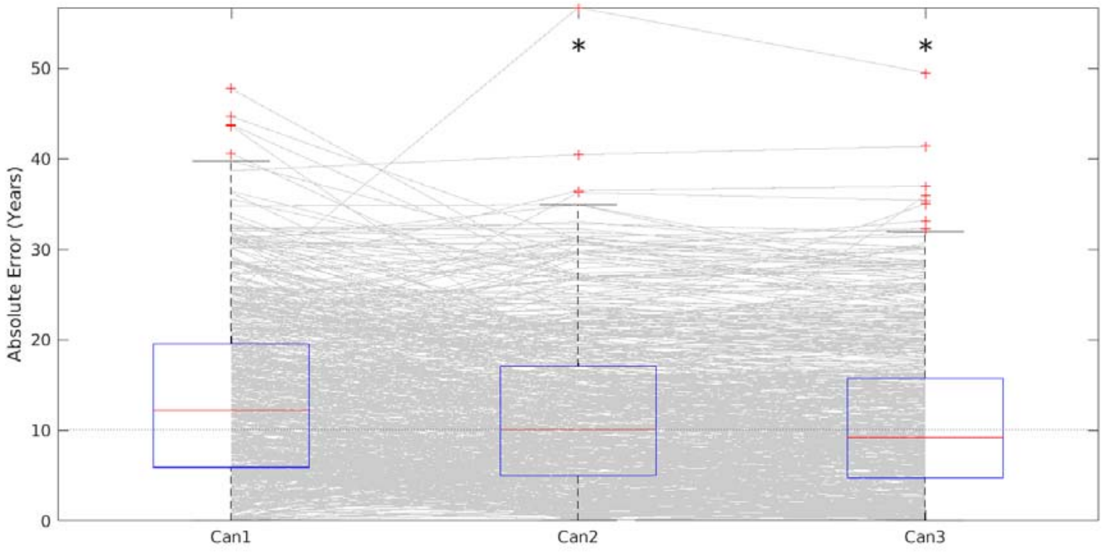


Supplementary Figure S6. Cross-validated absolute error in predicting age, using Can model parameters combined across all ROIs, as the number of basis functions increases, from canonical HRF only (Can1), to adding its temporal derivative (Can2) and then also its dispersion derivative (Can3). Each grey line corresponds to one participant. Superimposed on these lines are boxplots together with outliers (red crosses). An asterisk means that a sign-test revealed significantly better prediction than the simplest (Can1) model. The median error was 12.2 years (Can1), 10.0 years (Can2) and 9.2 years (Can3).

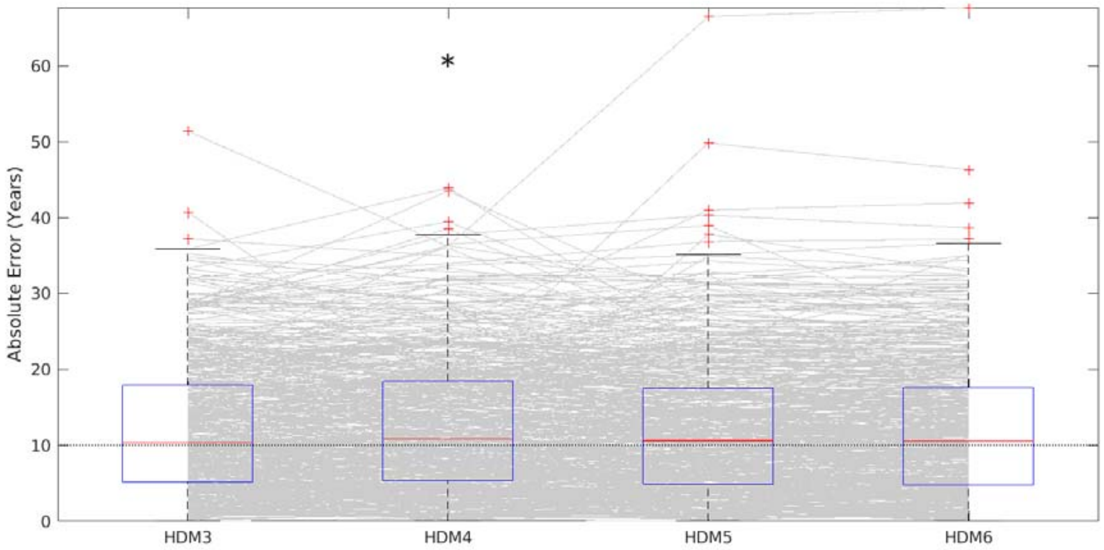


Supplementary Figure S7. Cross-validated absolute error in predicting age, using the HDM model parameters combined across all ROIs, as the number of parameters increased. The HDM3 model is as in main paper; an HDM4 model with the additional vessel stiffness parameter $\alpha$; an HDM5 model with the additional neurovascular feedback parameter $\gamma$; an HDM6 model with the additional oxygen extraction fraction $E_0$. Each grey line corresponds to one participant. Superimposed on these lines are boxplots together with outliers (red crosses). An asterisk means that a sign-test revealed significantly different prediction than the simplest (HDM3) model. The average across ROIs of the median error was 10.3 years (HDM3), 10.8 years (HDM4), 10.6 years (HDM5) and 10.5 years (HDM6). Note this is using the parameter estimates before application of PEB.

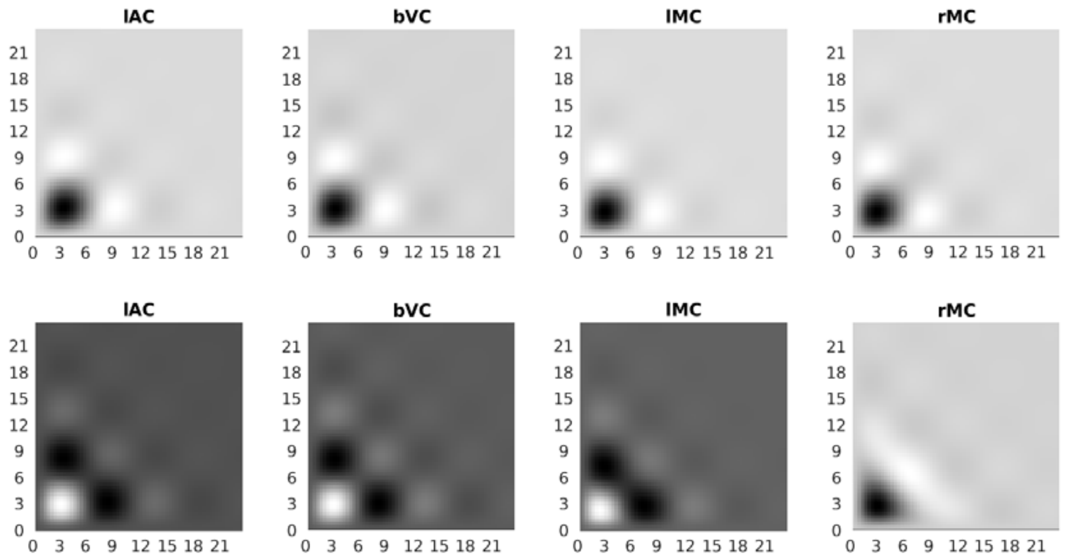


Supplementary Figure S8. Second-order Volterra kernels for each ROI from the HDM3 model, capturing nonlinear effects of time between trial onsets (SOA; plotted on both axes). The top row shows the mean across participants, while the bottom row shows the effect of age. Note that in order to see effects (e.g, in rMC), the grayscale is optimised for each ROI and kernel separately (units are arbitrary).
